# The Shifting Tempo of Evolution: Mapping Site-Specific Rate Shifts Across the Tree of Life

**DOI:** 10.64898/2026.06.19.733087

**Authors:** Muhammed Rasit Durak, Estelle Renaud, Julien Y. Dutheil

## Abstract

Evolutionary rates vary widely among sites in protein sequences, reflecting differences in functional constraints across residues. Individual sites can also experience lineage-specific shifts in substitution rates—a process known as heterotachy—when selective pressures change during evolution. Although such temporal variation has long been recognized, the mechanisms underlying lineage-specific rate shifts and the factors shaping their distribution across protein families remain poorly understood. Here we map site-specific rate shifts across thousands of orthologous protein families spanning the tree of life. Among more than 1.8 million aligned amino-acid sites, over one quarter show evidence of lineage-specific rate changes. Rate shifts are more frequent in families with deep evolutionary origins, including those tracing back to LUCA, whereas younger clade-specific families generally show lower proportions of rate-shifting sites. We next examined whether local structural features predict where rate shifts occur. Residue burial shows only a weak association with rate-shift probability, and its direction differs across domains, with buried residues enriched for rate shifts in Archaea but surface-exposed residues showing slightly higher probabilities in Eukaryota. Moreover, rate-shifting sites rarely form spatial clusters within protein structures, indicating that structural constraints do not globally determine their locations. Despite their widespread occurrence, rate-shifting sites have limited impact on phylogenetic reconstruction beyond random site variation. Together, these results show that lineage-specific rate shifts are a pervasive feature of protein evolution shaped primarily by evolutionary ancestry and phylogenetic depth.

## 1 Introduction

Proteins evolve under a shifting landscape of functional demands, environmental pressures, and lineage-specific histories (Yang 2014). Over deep evolutionary timescales, these forces leave heterogeneous signatures in molecular sequences, with some amino acid sites remaining highly conserved while others undergo bursts of rapid change (Penny, et al. 2001). Understanding how and why evolutionary rates vary across sites and across lineages is central to reconstructing the history of life, inferring adaptive processes, and accurately modeling molecular evolution. One particularly intriguing manifestation of evolutionary rate heterogeneity is the phenomenon whereby the evolutionary rate of a given site changes over time, a pattern known as heterotachy (Lopez, et al. 2002). Rather than evolving at a constant pace, individual residues may accelerate, decelerate, or alternate between conserved and variable states as selective pressures shift (Tuffley and Steel 1998). This temporal variation in site-specific evolutionary rates has long been recognized as a feature of molecular evolution, yet its prevalence and biological significance across the tree of life remain poorly understood. This gap in understanding is particularly important because patterns of evolutionary rate variation are not only of biological interest but also directly influence the assumptions and performance of phylogenetic models used to reconstruct evolutionary history.

Phylogenetic reconstruction lies at the heart of evolutionary biology, enabling inferences about species relationships, divergence times, and adaptive trajectories. At the molecular level, these inferences depend on substitution models—mathematical frameworks that balance biological realism with computational tractability (Liberles, et al. 2012; Sikosek and Chan 2014). Historically, substitution models have relied on simplifying assumptions to remain analytically tractable, even though these assumptions are known to be biologically imperfect (Liberles, et al. 2012). Early foundational models adopted particularly strong simplifications that nonetheless proved instrumental for the emergence of molecular phylogenetics. The Jukes–Cantor model, introduced in 1969 (Jukes and Cantor 1969), formalized sequence evolution under the assumption that all nucleotide substitutions occur with equal probability and at a constant rate across sites. Kimura’s two-parameter model subsequently relaxed this assumption by allowing different rates for transitions and transversions (Kimura 1980), yet still assumed uniform evolutionary rates along a sequence. As molecular data accumulated, it became increasingly clear that biological sequences violate assumptions of rate uniformity. Functional and structural constraints impose strong heterogeneity in evolutionary rates across sites (Worth, et al. 2009). Amino acid residues essential for maintaining protein structure or catalytic activity tend to be highly conserved, whereas residues located in flexible or solvent-exposed regions may tolerate greater variation (Smith, et al. 2007). These patterns reflect the underlying physical interactions among amino acids that stabilize protein structure and shape functional dynamics, indicating that evolutionary rates are tightly linked to site-specific structural context (Sikosek and Chan 2014). In particular, residues buried in the protein core tend to evolve more slowly than surface-exposed residues, which experience fewer structural constraints and greater evolutionary flexibility (Franzosa and Xia 2009). To accommodate this biological reality, models incorporating among-site rate variation (ASRV) were introduced in the 1990s, most notably through gamma-distributed rates or discrete rate categories (Yang 1994; Felsenstein and Churchill 1996). These models substantially improved realism and model fit by allowing different sites to evolve at different rates and are now standard in phylogenetic analyses. Despite their success, ASRV models retain a critical simplifying assumption: once assigned, a site’s evolutionary rate is assumed to remain constant throughout its evolutionary history (Yang 1996). Under this framework, a site evolving slowly in one lineage is expected to evolve slowly in all others, making site-specific rate constancy a cornerstone of most phylogenetic inference approaches. This assumption is at odds with biological intuition, as functional and structural constraints acting on individual residues are not static but can change over time due to functional divergence, environmental shifts, or compensatory mutations (Fitch 1971). If such changes alter the selective pressures acting on a site, they should be reflected by evolutionary rates that vary not only across sites but also across lineages (Yang 1996). The concept of heterotachy was introduced precisely to address this limitation. Its roots can be traced to the covarion model, first proposed in 1970, in which sites switch between variable (“on”) and conserved (“off”) states over time, with evolutionary changes occurring in a coordinated manner across sites rather than independently (Fitch and Markowitz 1970). Subsequent covarion-like models expanded this framework, permitting sites to accelerate, decelerate, or transition between evolutionary regimes along different branches of the tree (Tuffley and Steel 1998; Galtier 2001; Huelsenbeck 2002). These models explicitly acknowledged that selective pressures acting on sites may change through evolutionary time and provided a theoretical basis for capturing such dynamics within phylogenetic inference.

Yet, despite more than five decades of conceptual and methodological development, heterotachy has rarely been examined at scale in empirical datasets, particularly in conjunction with protein structural information. Most large phylogenomic analyses continue to rely on ASRV-based frameworks, in part due to their computational efficiency and the limited availability of accessible heterotachy-aware implementations. As a consequence, it remains unclear how prevalent heterotachy truly is across the tree of life, whether it is restricted to particular genes or lineages, or how it interacts with structural constraints that are known to shape evolutionary rates across sites. This gap has important implications: if heterotachy is widespread, failure to account for lineage-specific rate shifts may systematically distort phylogenetic reconstruction, molecular dating, and ancestral state inference, particularly when convergent evolution, homoplasy, or episodic selection are misinterpreted as consistent phylogenetic signal. These unresolved issues motivate two fundamental questions in molecular evolution: (1) Is heterotachy a rare, lineage-specific phenomenon, or a widespread and intrinsic feature of protein evolution? (2) What factors underlie site-specific rate shifts—do they correlate with protein structural features, lineage-specific adaptations, or reflect largely stochastic evolutionary processes? In this study, we refer to lineage-specific changes in site-specific evolutionary rates as rate shifts, using this term operationally to describe empirical instances of heterotachy inferred from sequence data. To address these questions, we perform a large-scale, phylogenetically comprehensive analysis of rate shifts across the tree of life. Using orthologous protein sequences from over 2,900 species spanning all three domains of life, we infer site-specific evolutionary rate shifts and assess their distribution across diverse taxonomic lineages. Beyond quantifying the prevalence of rate shifts, we integrate structural bioinformatics by mapping rate-shifting sites onto experimentally determined protein structures to test whether rate shifts preferentially occur in specific structural contexts, such as solvent-exposed regions, active sites, or interaction interfaces. Together, our study characterizes the global landscape of heterotachy, identifies its evolutionary correlates, and clarifies its methodological implications, providing an empirical foundation for models of sequence evolution that more accurately reflect the dynamic nature of molecular change.

## 2 Results

### 2.1 Large-scale orthologous protein dataset

Our dataset comprises 6,012 orthologous protein families obtained from the OMA database (Altenhoff, et al. 2024). Individual orthogroups contain between 100 and 2,670 protein sequences, for a total of approximately 2.4 million sequences across all families. The number of sequences per orthogroup varies widely but is concentrated around a few hundred species per family (Supplementary Figure S1). After alignment trimming and quality filtering, cleaned multiple sequence alignments range from 49 to 4,653 amino-acid positions. Across all orthogroups this corresponds to approximately 2.6 million high-confidence alignment sites. Rate-shift inference using The RAte Shift EstimatoR (RASER) (Penn, et al. 2008) successfully converged for 4,125 orthologous groups. These families include 1,816,880 aligned amino-acid sites that were used in all subsequent analyses. Each of the 4,125 orthologous groups was classified according to its inferred last common ancestor (LCA), resulting in 29 distinct age categories. Orthogroups are distributed across both deep and more recently derived evolutionary nodes (Figure 1a). A large fraction of families trace back to deep ancestral clades, including 888 orthogroups assigned to the Last Universal Common Ancestor (LUCA) and 1,210 assigned to Eukaryota (Figure 1a). Additional groups originate within major domains such as Bacteria (253) and Archaea (23) (Figure 1a). Within eukaryotes, orthogroups are further distributed across multiple nested clades—including Bilateria, Eumetazoa, and Gnathostomata—with smaller numbers associated with more specific vertebrate and chordate lineages. Overall, the dataset spans a broad range of evolutionary depths, from ancient protein families conserved across domains of life to lineage-restricted orthogroups emerging within more recent eukaryotic clades.

**Figure 1.**
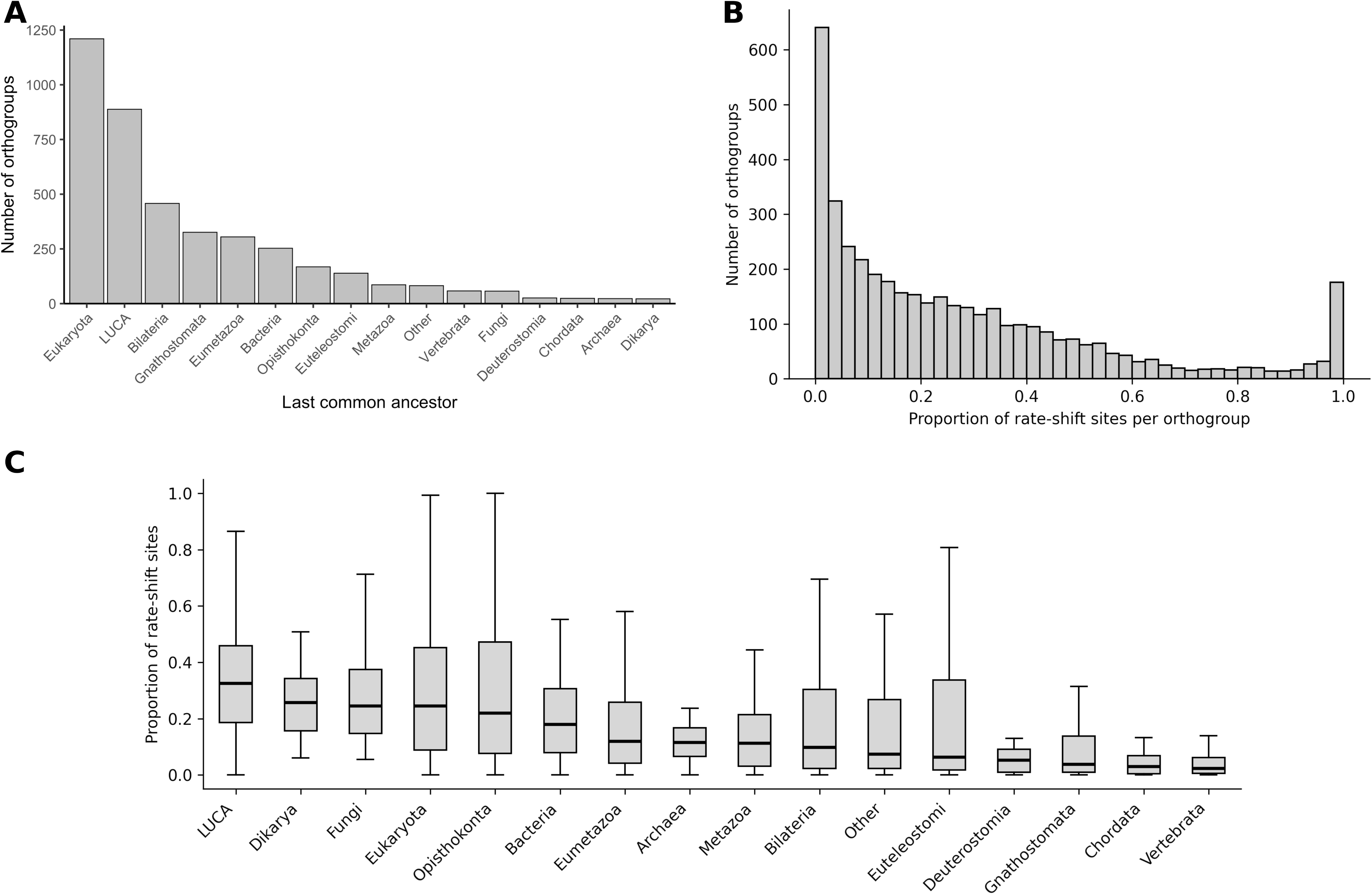
Distribution of orthogroups and rate-shift prevalence across evolutionary ancestry. (A) Number of orthogroups assigned to each inferred last common ancestor (LCA) category, with only the 15 most frequent categories shown explicitly and all remaining grouped as “Other”. (B) Distribution of the proportion of rate-shifting sites across orthogroups, with most families exhibiting relatively low proportions but substantial variation observed across the dataset. (C) Distribution of rate-shift proportions across orthogroups grouped by their inferred last common ancestor (LCA), with categories ordered by decreasing median rate-shift proportion and less frequent categories grouped as “Other”.

### 2.2 Site-specific rate shifts are widespread across protein families

Among the 1,816,880 aligned amino-acid sites analyzed, 504,232 posterior probabilities greater than 0.95 and were therefore classified as rate-shifting, corresponding to 27.8% of all positions tested. The median proportion per family was 0.19, meaning that half of the 4,125 orthogroups contained rate shifts at 19% or fewer of their aligned sites. However, numerous families exhibited considerably higher fractions, with rate-shifting sites affecting a large share of positions within some orthogroups (Figure 1b). Of the 1,816,880 total aligned sites, 263,483 could be mapped to experimentally determined protein structures. This structurally resolved subset included both rate-shifting and non–rate-shifting sites and was used for downstream structural analyses. When orthogroups were grouped according to their inferred last common ancestor (LCA), differences in rate-shift prevalence became apparent across evolutionary ancestry categories (Figure 1c).

When orthogroups were grouped according to their inferred last common ancestor (LCA), rate-shift proportions differed significantly among categories (Kruskal–Wallis test, H = 536.8, df = 15, p < 0.001), with Dunn’s post hoc tests confirming significant pairwise differences among major evolutionary groups. Orthogroups originating at deep evolutionary levels, particularly those assigned to LUCA and broad eukaryotic categories, tended to exhibit higher proportions of rate-shifting sites. This pattern is also observed at the structural level, with representative protein families spanning different evolutionary depths showing variation in the distribution of rate-shifting sites (Figure 2). In contrast, more recently derived and nested clades, including vertebrate and chordate categories, generally showed lower heterotachy prevalence (Supplementary Figure S2 and Table S1).

**Figure 2.**
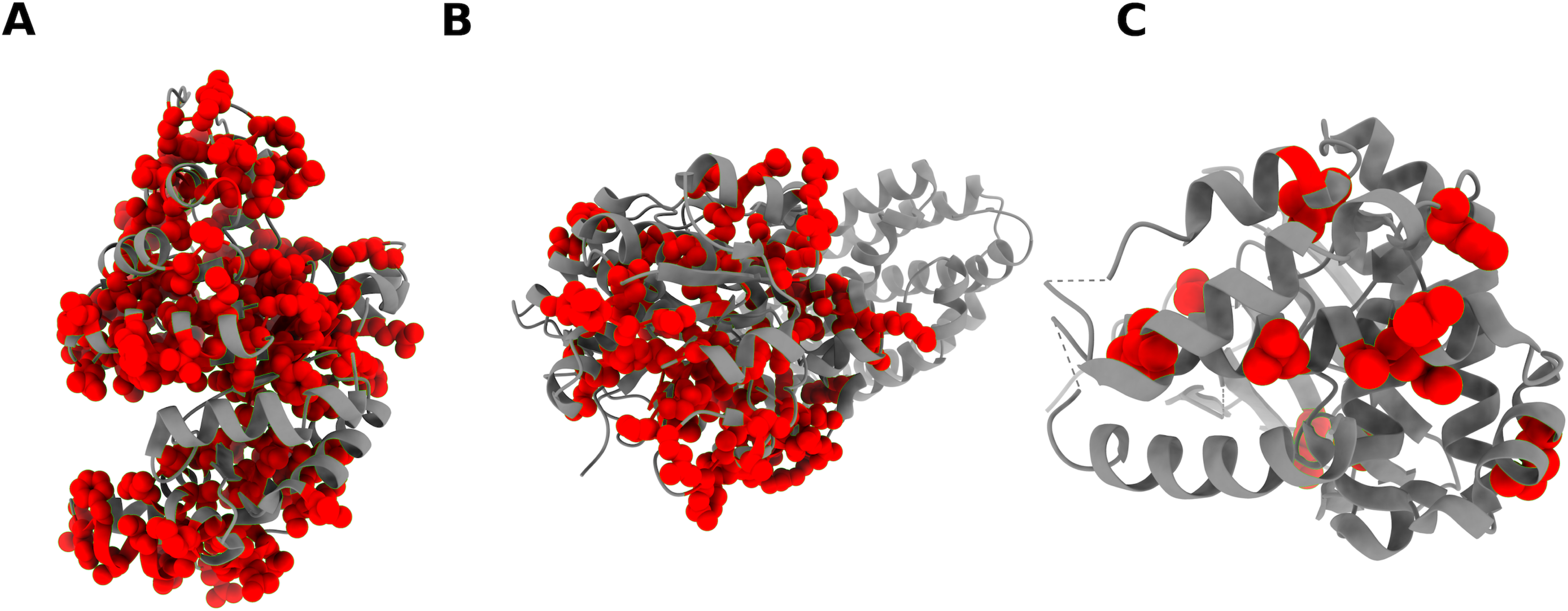
Representative protein structures illustrating variation in the distribution of rate-shifting sites across protein families with different evolutionary ancestries. Shown are examples of families inferred to originate from (A) LUCA (PDB: 1EZW), (B) Bacteria (PDB: 6JZU), and (C) Bacilli (PDB: 7S3L). Rate-shifting sites are highlighted in red, illustrating differences in their distribution across families spanning deep to more recent evolutionary origins.

### 2.3 Evolutionary ancestry and phylogenetic scale are associated with family-level rate-shift prevalence

To identify factors associated with variation in the prevalence of rate shifts among protein families, we tested whether phylogenetic scale (total tree length and number of species/sequences per orthogroup) and evolutionary ancestry (inferred last common ancestor, used as a proxy for evolutionary age) predict differences in the proportion of rate-shifting sites among orthogroups. Family-level heterotachy prevalence showed significant positive associations with both total phylogenetic tree length (Kendall’s τ = 0.38, p < 0.001) and the number of sequences per orthogroup (τ = 0.32, p < 0.001), indicating that orthogroups spanning greater evolutionary divergence and broader taxonomic sampling tend to exhibit higher proportions of rate-shifting sites. Because deeply conserved families may also be more broadly sampled and span longer evolutionary timescales, we next performed residual-based partial correlation analyses to evaluate the independent contributions of these factors. After correcting for LCA category, the number of sequences per orthogroup remained positively associated with the proportion of rate-shifting sites (τ = 0.26, p < 0.001), indicating that broader taxonomic sampling independently contributes to the prevalence or detectability of rate shifts. Similarly, total phylogenetic tree length remained significantly associated with heterotachy prevalence after correcting for both LCA category and the number of sequences per orthogroup (τ = 0.17, p < 0.001), suggesting that overall sequence divergence also contributes independently to lineage-specific rate variation.

### 2.4 Structural burial shows only a small effect relative to sampling and divergence

We next evaluated whether local structural features predict the probability that an alignment site is classified as rate-shifting, while accounting for variation in evolutionary divergence and taxonomic sampling. We fitted a site-level mixed-effects logistic regression model including structural predictors (relative solvent accessibility (RSA), Cα depth, and secondary structure), phylogenetic covariates (tree length and number of sequences per orthogroup), and orthogroup identity as a random effect (Table 1; Supplementary Table S2). RSA quantifies residue solvent exposure, whereas Cα depth more directly captures structural burial.

**Table 1.**
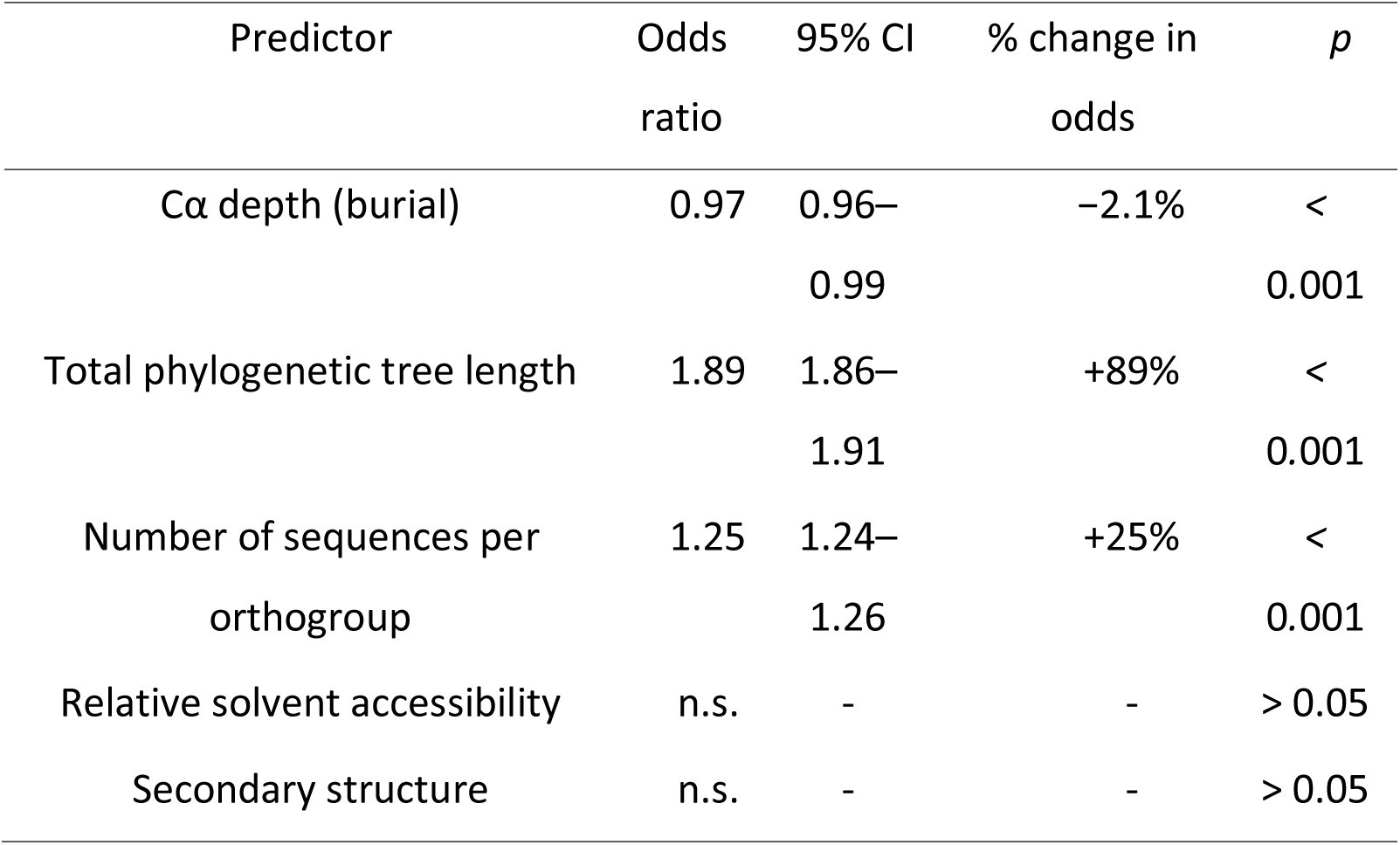
Baseline mixed-effects logistic regression model predicting the probability that an alignment site is classified as rate-shifting. Continuous predictors were standardized (mean = 0, SD = 1), so odds ratios represent the effect of a one standard deviation increase in each variable. Percent change in odds was calculated as (OR − 1) × 100. Odds ratios below 1 indicate reduced probability of rate shifts, whereas values above 1 indicate increased probability. n.s. denotes predictors that were not statistically significant (*p >* 0.05).

Among the structural predictors included in the baseline model, residue burial measured as Cα depth was the only variable showing a statistically significant association with rate-shift probability (odds ratio (OR) = 0.97 per standard deviation increase in depth, 95% CI: 0.96–0.99; p < 0.001; Table 1), whereas RSA and secondary structure class showed no significant effects. An odds ratio below one indicates that more deeply buried residues are less likely to be rate-shifting. Specifically, a one-standard-deviation increase in burial corresponds to an approximately 2.1% decrease in the odds that a site is classified as rate-shifting. Thus, surface-exposed residues show a modestly higher probability of rate shifts. Despite statistical significance, the magnitude of this structural effect was small. In contrast, variables reflecting evolutionary divergence and taxonomic sampling exhibited substantially larger associations. Total phylogenetic tree length was associated with an 89% increase in the odds of detecting a rate shift per standard deviation increase (OR = 1.89), and the number of sequences per orthogroup with a 25% increase (OR = 1.25; both p < 0.001).

### 2.5 Evolutionary age explains substantial baseline differences in rate-shift probability

We next incorporated evolutionary ancestry as a four-level categorical predictor (Archaea, Bacteria, Eukaryota, and LUCA-derived families) to evaluate whether baseline probabilities of rate-shifting sites differ among major evolutionary groups. Results from this mixed-effects model are summarized in Table 2, with the full model output reported in Supplementary Table S3. After including ancestry, marked differences in baseline odds were observed across domains. Relative to Archaea, Bacteria showed a 33% increase in the odds that a site is classified as rate-shifting (OR = 1.33), LUCA-derived families showed a 129% increase (OR = 2.29), and Eukaryota exhibited a 204% increase (OR = 3.04). All pairwise comparisons were statistically significant. Importantly, inclusion of ancestry did not materially change the structural coefficient for residue burial. Depth remained negatively associated with rate-shift probability (OR = 0.980 per 1 standard deviation increase), corresponding to an approximate 2% decrease in the odds of a site being classified as rate-shifting. Relative solvent accessibility and secondary structure remained non-significant. Covariates reflecting evolutionary divergence and sampling retained strong effects: total phylogenetic tree length was associated with more than a two-fold increase in odds per standard deviation (OR = 2.17), and the number of sequences per orthogroup with a 20% increase (OR = 1.20). Overall, baseline differences among major evolutionary domains substantially exceeded the magnitude of local structural effects.

**Table 2.**
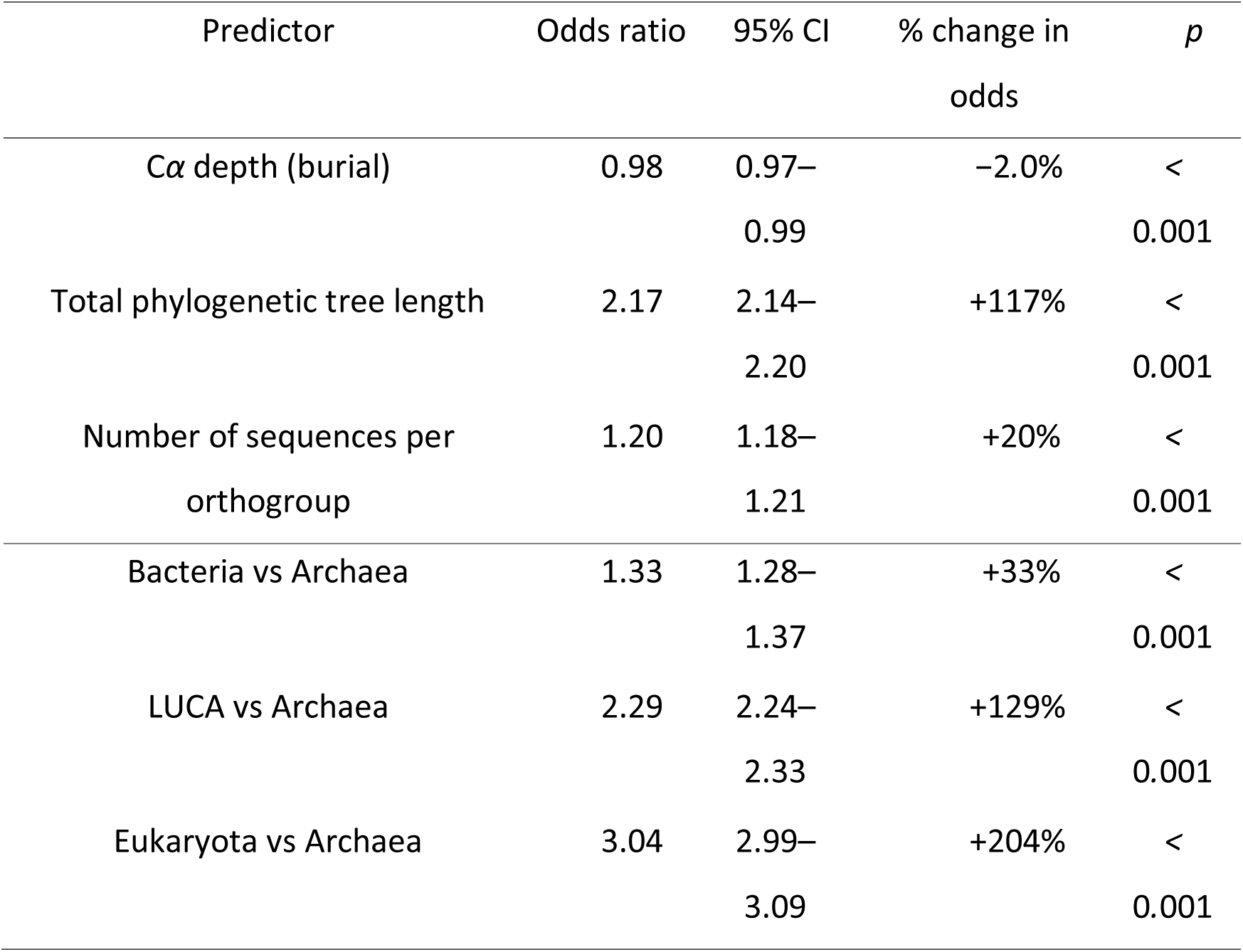
Mixed-effects logistic regression model including evolutionary ancestry as a categorical predictor of rate-shift probability. Continuous predictors were standardized (mean = 0, SD = 1), and odds ratios therefore represent the effect of a one standard deviation increase in each continuous variable. Percent change in odds was calculated as (OR − 1) × 100. Odds ratios greater than 1 indicate higher odds of rate-shifts relative to the reference category (Archaea).

| Predictor | Odds ratio | 95% CI | % change in odds | *p* |
| --- | --- | --- | --- | --- |
| *Cα* depth (burial) | 0.98 | 0.97–<br>0.99 | –2.0% | <<br>0.001 |
| Total phylogenetic tree length | 2.17 | 2.14–<br>2.20 | +117% | <<br>0.001 |
| Number of sequences per orthogroup | 1.20 | 1.18–<br>1.21 | +20% | <<br>0.001 |
| Bacteria vs Archaea | 1.33 | 1.28–<br>1.37 | +33% | <<br>0.001 |
| LUCA vs Archaea | 2.29 | 2.24–<br>2.33 | +129% | <<br>0.001 |
| Eukaryota vs Archaea | 3.04 | 2.99–<br>3.09 | +204% | <<br>0.001 |

### 2.6 The effect of structural burial differs across evolutionary domains

Because residue burial showed a significant association with rate-shift probability in previous models, we next tested whether the effect of burial varies across major evolutionary domains by including an interaction term between C*α* depth and evolutionary ancestry. Results from this interaction model are summarized in Table 3, with the full model results provided in Supplementary Table S4. Within the reference category (Archaea), burial showed a positive association with rate-shift probability (OR *>* 1; Table 3), indicating that more deeply buried residues in archaeal proteins exhibit higher odds of being classified as rate-shifting. Thus, in archaeal proteins, core residues exhibit a higher probability of rate shifts. Interaction terms quantify how this burial effect differs across other evolutionary domains relative to Archaea. For Eukaryota, the depth interaction term was negative and statistically significant, reversing the direction of the archaeal effect. In eukaryotic families, increasing burial was associated with reduced odds of rate shifts, indicating enrichment of rate shifts at more surface-exposed residues. In contrast, interaction terms for Bacteria and LUCA-derived families were not significant, indicating that solvent exposure does not exert a strong or consistent influence on rate-shift probability in these groups. Overall, these results suggest that local structural features are not globally consistent predictors of rate-shifting sites across evolutionary lineages. While archaeal proteins show an enrichment of rate shifts in more buried residues and eukaryotic proteins show a modest enrichment toward surface-exposed sites, no consistent structural bias is observed in bacterial or LUCA-derived families.

**Table 3.** Mixed-effects logistic regression model testing whether the effect of residue burial (C*α* depth) on rate-shift probability differs across evolutionary domains. Archaea was used as the reference category. Interaction terms represent the change in the burial effect relative to Archaea. Continuous predictors were standardized (mean = 0, SD = 1). Percent change in odds was calculated as (OR−1)×100.

| Predictor | Odds ratio | 95% CI | % change in odds | *p* |
| --- | --- | --- | --- | --- |
| $C\alpha$ depth (Archaea baseline) | 1.46 | 1.44–1.48 | +46.0% | < 0.001 |
| Depth × Bacteria | 0.65 | 0.62–0.68 | –34.8% | < 0.001 |
| Depth × Eukaryota | 0.67 | 0.65–0.68 | –33.3% | < 0.001 |
| Depth × LUCA | 0.68 | 0.67–0.69 | –31.9% | < 0.001 |

To further examine lineage-specific structural effects, we fitted separate mixed-effects models within each major evolutionary domain: Archaea (1,991 sites across 23 families), Bacteria (24,834 sites across 235 families), Eukaryota (176,426 sites across 1,705 families), and LUCA-derived families (54,189 sites across 649 families). These lineage-specific models retained the same structural predictors, phylogenetic covariates, and orthogroup random intercept as the global models, allowing structural associations to vary among evolutionary domains. The full results of these domain-specific models are provided in Supplementary Tables S5–S8. Within Archaea, increasing burial was associated with a 22% increase in the odds of a site being classified as rate-shifting per standard deviation increase in depth, consistent with the depth-by-ancestry interaction observed in the global model (Figure 3). This pattern indicates an enrichment of rate shifts among more deeply buried residues in archaeal proteins. In contrast, in Eukaryota, increasing burial was associated with an approximately 3% decrease in odds, corresponding to a modest enrichment of rate shifts at more surface-exposed residues. In Bacteria and LUCA-derived families, burial was not significantly associated with the probability of rate shift (Figure 3). Taken together, these results indicate that the structural burial effect observed in the global model does not reflect a uniform pattern across evolutionary domains. Instead, it arises from contrasting trends in Archaea and Eukaryota, with no detectable structural bias observed in Bacteria or in LUCA-derived families. Across all domains, covariates capturing evolutionary divergence and sequence sampling showed substantially larger positive associations with rate-shift probability, and their magnitudes exceeded those of structural predictors. These might highlight pronounced differences among evolutionary domains and indicate that variation in evolutionary divergence and taxonomic sampling contributes more strongly to the prevalence of rate shifts than local structural features.

**Figure 3.**
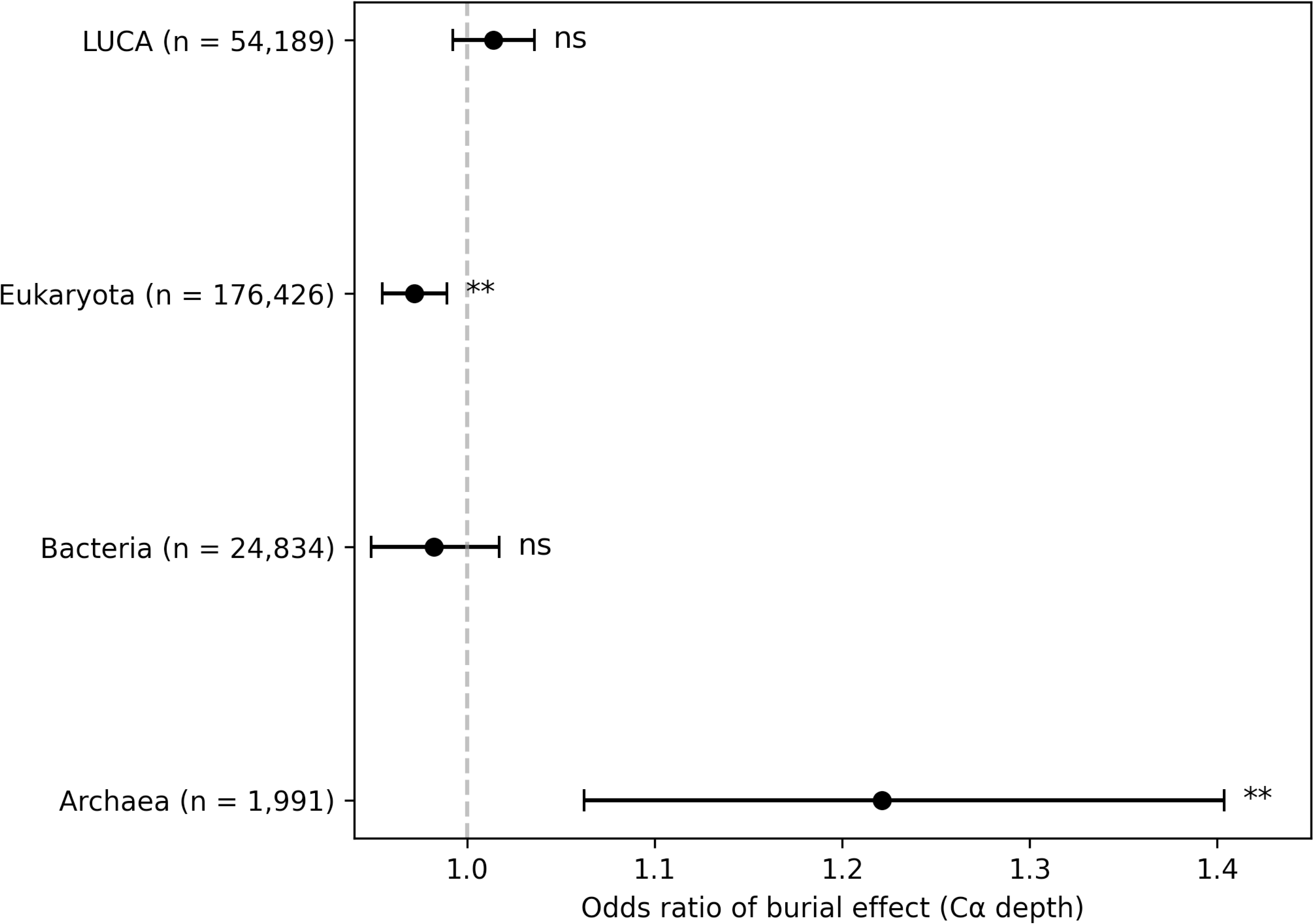
Domain-specific effects of residue burial on the probability that an alignment site is classified as rate-shifting. Odds ratios and 95% confidence intervals for the effect of C*α* depth were estimated from mixed-effects models fitted separately within Archaea, Bacteria, LUCA-derived families, and Eukaryota. Sample sizes (n) correspond to the number of structurally mapped sites included in each model. The dashed vertical line indicates an odds ratio of 1 (no effect). Asterisks denote statistically significant effects (*p <* 0.05).

### 2.7 Rate-shifting sites are rarely clustered within protein structures

To evaluate whether rate-shifting sites tend to occur in spatially localized regions of protein structures, we analyzed their three-dimensional proximity across 1,906 protein families containing at least two structurally mappable sites. If rate-shifting sites preferentially accumulate in specific structural regions, they would be expected to form spatial clusters within protein structures. The distribution of permutation-based *p*-values was broadly consistent with random expectation showing no widespread tendency for rate-shifting sites to occur in close three-dimensional proximity (Supplementary Figure S3). After correction for multiple testing using the Benjamini–Hochberg false discovery rate procedure (q ≤ 0.05), significant spatial clustering was detected in only 76 families (4.0%), a proportion below the 5% expected by chance under the null hypothesis, whereas the large majority of proteins showed dispersion patterns indistinguishable from random expectation. In most proteins, rate-shifting sites were distributed across different regions of the structure rather than forming localized structural clusters, indicating that spatial aggregation of rate shifts is potentially limited to a small subset of protein families.

### 2.8 The impact of rate-shifting sites on phylogenetic reconstruction is limited

Across 50 gene families, the removal of rate-shifting (RS) sites resulted in only small changes in tree topology. Differences in normalized Robinson–Foulds distances (nRF) between RS-removed trees and randomly subsampled trees, calculated relative to the corresponding full-alignment tree, were generally small, with many gene families showing slightly positive shifts, although values remained close to zero overall (Figure 4A). Positive values indicate that removal of RS sites produced larger topological differences from the full-alignment tree than expected under random site removal, whereas negative values indicate the opposite. While some gene families showed larger topological differences after RS-site removal, these effects were not systematically greater than those observed under random site removal. Permutation tests performed for each gene family showed that only 5 out of 50 families (10%) exhibited significant differences (p < 0.05), indicating that, in most cases, the topological impact of removing RS sites is comparable to that expected from random removal of sites. Variation in total phylogenetic tree length among random-removal replicates was generally centered around the orthogroup-specific mean, although the magnitude of variation differed substantially among gene families (Figure 4B). Most distributions were tightly clustered around zero, indicating that random site removal had relatively limited effects on overall branch length estimation, while a small number of orthogroups exhibited broader distributions and occasional extreme values. A similar pattern was observed for mean bootstrap support (Figure 4C). Support values from random-removal replicates were centered on the orthogroup-specific mean and showed relatively narrow distributions for most gene families, although several orthogroups displayed greater variability. Overall, these results indicate that random removal of sites introduces only modest variation in branch support across the majority of gene families. Linear models were used to test whether variation in the phylogenetic effects of RS-site removal across gene families could be explained by orthogroup characteristics. Neither the number of sequences, alignment length, nor the proportion of rate-shifting sites showed significant associations with differences in nRF topology scores (all p > 0.23), branch-length effects (all p > 0.33), or bootstrap-support effects (all p > 0.09), indicating that variation in the phylogenetic impact of RS-site removal is not explained by these orthogroup properties.

**Figure 4.**
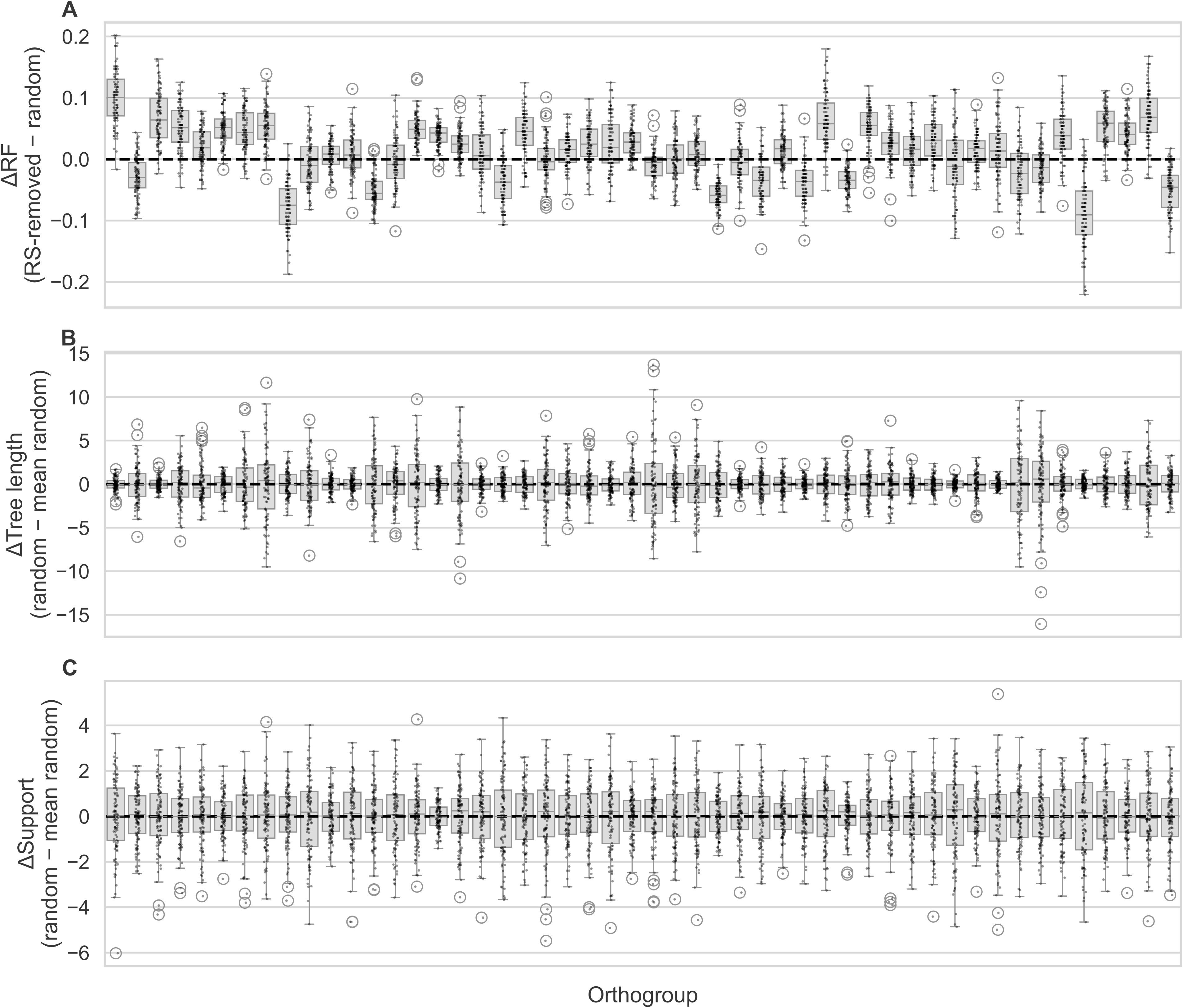
Effects of removing rate-shifting sites on phylogenetic topology, tree length, and branch support across 50 gene families. (A) Differences in normalized Robinson–Foulds distances (ΔRF; RS-removed − random). (B) Differences in tree length relative to the mean of random-removal replicates (ΔTree length; random − mean random). (C) Differences in mean branch support relative to the mean of random-removal replicates (ΔSupport; random − mean random). Boxes represent interquartile ranges, horizontal lines indicate medians, and points correspond to individual replicate analyses. In panel A, the baseline is the value obtained after random site removal for the same replicate. In panels B and C, the baseline is the mean value across all random-removal replicates within each orthogroup. Positive values indicate observations above the respective baseline, whereas negative values indicate observations below the baseline.

## 3 Discussion

Our large-scale study across 4,125 orthologous protein families spanning the tree of life demonstrates that lineage-specific shifts in site-level evolutionary rates are widespread. These findings indicate that temporal variation in evolutionary constraints at individual sites is a general feature of protein evolution rather than a marginal phenomenon, consistent with early theoretical expectations of rate variation through time (Fitch 1971). The possibility that evolutionary rates may change over time at individual sites has long been recognized in theoretical models of sequence evolution. Despite these conceptual advances, empirical quantification of the prevalence and distribution of such temporal rate variation across large numbers of protein families has remained limited. Our analysis helps bridge this gap by providing large-scale empirical evidence that heterotachy is pervasive throughout diverse evolutionary lineages. Rate shifts were not evenly distributed across protein families. Orthogroups inferred to have originated at deep ancestral nodes (e.g., those tracing back to LUCA) tended to exhibit higher proportions of rate-shifting sites than more recently derived clades. This suggests that lineage-specific changes in site-specific constraints accumulate over evolutionary time. Ancient protein families have experienced prolonged exposure to changing cellular environments, lineage-specific functional divergence, and shifting interaction networks, which may repeatedly remodel the selective pressures acting on individual residues and produce episodes of rate acceleration or deceleration across lineages (Fitch 1971). In contrast, younger protein families restricted to narrower phylogenetic ranges have had fewer opportunities for such constraint reorganization. Associations between heterotachy prevalence and phylogenetic scale further support this interpretation. Orthogroups characterized by greater evolutionary divergence and broader sequence representation tended to exhibit higher proportions of rate-shifting sites.

These findings also highlight a broader conceptual point about the nature of molecular evolution. Protein sequences are often interpreted through the lens of relatively stable structural constraints, where individual residues occupy fixed positions within a landscape of functional and biophysical requirements (Franzosa and Xia 2009; Sikosek and Chan 2014). However, our results suggest that the selective environment experienced by individual sites is inherently dynamic over evolutionary time. As proteins adapt to new cellular contexts, interaction partners, and environmental conditions, the functional importance of specific residues may shift, leading to episodic changes in evolutionary rate. In this view, heterotachy emerges not as an anomalous feature of sequence evolution but as a natural consequence of evolving biological systems whose functional constraints continually change across lineages.

In contrast to the strong influence of evolutionary ancestry and phylogenetic scale, local structural features exerted comparatively modest effects. Residue burial showed only a small association with rate-shift probability in the global analyses, while relative solvent accessibility and secondary structure class showed no independent effects after accounting for ancestry and phylogenetic opportunity. Moreover, rate-shifting sites were rarely spatially clustered within protein structures, with significant clustering detected in only a small minority of protein families. Together, these observations indicate that structural features do not exert a global constraint on the locations of rate shifts. Instead, structural influences appear to operate in a lineage-dependent manner. Domain-specific analyses revealed contrasting patterns between major evolutionary groups. In particular, archaeal proteins showed an enrichment of rate shifts among more deeply buried residues, whereas eukaryotic proteins exhibited a modest enrichment toward surface-exposed sites. Such differences suggest that structural determinants of evolutionary change may reflect lineage-specific selective pressures rather than universal constraints.

The enrichment of rate shifts within buried residues in Archaea may be consistent with known patterns of protein adaptation in archaeal extremophiles. Many archaeal species inhabit environments characterized by extreme temperature, salinity, or acidity, conditions that impose strong stability constraints on protein structures (Vieille and Zeikus 2001; Reed, et al. 2013). Under such conditions, the hydrophobic core plays a critical role in maintaining structural integrity, and adaptive modifications often involve remodeling of core residues to optimize packing interactions and stabilize protein folding (Kumar and Nussinov 2001; Reed, et al. 2013). Consistent with this interpretation, several archaeal protein families in our dataset correspond to core informational proteins involved in transcription, translation, and DNA replication, including ribosomal proteins, transcription factors, and DNA replication enzymes (Supplementary Table S9). These proteins frequently form large macromolecular complexes and operate under strong structural constraints, which may favor adaptive remodeling of buried residues to maintain stability and functional interactions under extreme environmental conditions. One possible explanation is that adaptation to contrasting environmental conditions repeatedly alters the stability requirements of archaeal proteins. Because buried residues make major contributions to protein folding and thermodynamic stability, changes in temperature, salinity, or pH may require compensatory adjustments within the protein core to maintain function. Under this scenario, residues that are strongly constrained in one lineage may become targets of adaptive change in another lineage experiencing different environmental conditions, leading to shifts in evolutionary rate through time. Recurrent transitions among distinct ecological niches could therefore generate elevated levels of heterotachy within structurally buried regions even when overall structural constraints remain strong. In addition, archaeal proteins often rely on finely balanced trade-offs between stability and conformational dynamics, particularly under extreme conditions. Changes in the optimal balance between rigidity and flexibility across lineages may further alter the selective pressures acting on core residues, contributing to temporal variation in site-specific evolutionary rates. This contrasts with the typical pattern observed in many bacterial and eukaryotic proteins, where core residues tend to evolve more slowly due to strong structural constraints that limit acceptable substitutions (Franzosa and Xia 2009; Sikosek and Chan 2014). In the baseline mixed-effects model fitted across all sites, residue burial showed a small negative association with rate-shift probability, suggesting that more deeply buried residues were slightly less likely to experience rate shifts. Because this global model estimates a single structural effect across all evolutionary groups, the resulting coefficient reflects an average slope across heterogeneous lineages (Gelman and Hill 2007). When structural effects differ in direction among groups, such averaging can reduce the magnitude of the overall estimate and potentially obscure lineage-specific patterns. Consistent with this interpretation, the interaction model revealed that the weak global association between burial and rate-shift probability largely reflects the combination of contrasting domain-specific trends. Despite its limited representation, archaeal families consistently showed a positive association between residue burial and rate-shift probability, suggesting that lineage-specific structural adaptation may play a more prominent role in this domain. More broadly, the absence of strong global structural signatures suggests that heterotachy frequently arises from lineage-specific evolutionary processes rather than universal structural determinants. Rate shifts may reflect episodes of functional divergence, compensatory substitutions, or changes in molecular interaction networks that alter selective pressures on individual residues (Pazos and Valencia 2008). Such processes may operate differently across evolutionary lineages and over long timescales, producing heterogeneous patterns of constraint remodeling across protein families.

The pervasive nature of lineage-specific rate shifts also raises important considerations for phylogenetic inference. To directly evaluate whether rate-shifting sites influence phylogenetic inference, we performed a controlled site-removal analysis across 50 gene families. Removal of rate-shifting sites resulted in only small changes in tree topology, with differences in normalized Robinson-Foulds distances generally remaining close to those observed under random site removal. Although some gene families exhibited larger shifts, these effects were not systematic and, in most cases, were not statistically distinguishable from random expectations. These findings suggest that, despite the widespread presence of heterotachy, its impact on phylogenetic reconstruction may be limited because rate-shifting events are distributed across both sites and evolutionary lineages rather than concentrated in a manner that consistently favors alternative phylogenetic relationships. While lineage-specific rate shifts reflect dynamic changes in selective constraints at individual residues, their effects may be dispersed across different branches of the phylogeny, reducing the likelihood of generating systematic topological biases. Consequently, the phylogenetic signal contributed by rate-shifting sites may not act coherently in a single direction, helping to explain the limited effects observed following their removal. In this sense, heterotachy appears to be an intrinsic feature of protein evolution without necessarily introducing strong or systematic biases in phylogenetic reconstruction under standard modeling approaches. Many widely used substitution models incorporate variation in evolutionary rates among sites but assume that these rates remain constant through time (Yang 1994). Our results suggest that this assumption may oversimplify long-term patterns of molecular evolution, particularly across deep phylogenetic divergences where lineage-specific shifts in constraint are common. Models allowing temporal variation in site-specific constraints, such as covarion-like frameworks (Fitch and Markowitz 1970; Tuffley and Steel 1998), may therefore provide a more realistic representation of protein evolution over extended evolutionary timescales. Despite the large scale of the dataset, several limitations should be considered when interpreting these results. Structural analyses were restricted to sites with available experimentally determined structures, representing a subset of aligned positions. Inference of rate shifts depends on model assumptions and posterior probability thresholds, and detection power varies with phylogenetic depth and sampling density. The impact of rate-shifting sites on the phylogenetic reconstruction was restricted to a subset of 50 gene families, although each required extensive phylogenetic reconstruction across multiple replicate alignments. In addition, phylogenetic support was assessed using ultrafast bootstrap approximation, which may not fully capture uncertainty in all cases. Taken together, these results support the view that heterotachy is an intrinsic feature of protein evolution whose prevalence increases with evolutionary depth. Lineage history and evolutionary time appear to play dominant roles in shaping patterns of rate variation across sites, whereas structural constraints modulate these dynamics in lineage-specific ways rather than imposing universal restrictions. This perspective highlights the importance of considering both evolutionary history and structural context when interpreting patterns of molecular evolution across the tree of life.

## 4 Methods

### 4.1 Orthologous Sequence Database Construction

We used the July 2024 release of the OMA (Orthologous MAtrix) database as the foundation for constructing our set of orthologous protein groups (Altenhoff, et al. 2024). OMA groups consist of protein sequences inferred to be related exclusively through speciation and are free from duplication events. We first downloaded the complete set of OMA groups, comprising approximately 1.4 million groups spanning more than 2,900 species. To ensure sufficient sequence representation for robust phylogenomic analyses and adequate statistical power in downstream analyses, we applied an initial filtering criterion retaining only groups containing at least 100 sequences. This step reduced the dataset to 27,304 orthologous groups. We then further refined the dataset by selecting only those groups that included at least one protein with an experimentally determined three-dimensional structure. Structural information was obtained by cross-referencing OMA sequence identifiers with UniProt entries. After this final filtering step, a total of 6,012 orthologous groups were retained for downstream analyses. For each of the 6,012 retained groups, we inferred the last common ancestor (LCA) to represent the evolutionary age of the group. LCA assignment was performed using the getOMAGroup and getTaxonomy functions provided by the OmaDB R package (Kaleb, et al. 2019).

### 4.2 Alignment of Orthologous Groups

The 6,012 orthologous groups of protein sequences were aligned using two multiple sequence alignment tools: MUSCLE5 with the Super5 algorithm (Edgar 2021), which is optimized for speed and scalability on large datasets, and MAFFT version 7.3 (Katoh and Standley 2013). To evaluate and filter alignment quality, we employed the column score (CS) metric as defined by (Thompson, et al. 1999), which assesses whether corresponding alignment columns are identical between alternative alignments. Based on CS evaluation, a consensus alignment was generated for each orthologous group. Consensus alignments from both aligners were used for all downstream analyses. Additional quality control included manual inspection of a subset of alignments to verify overall consistency and alignment accuracy. Alignments were further filtered to retain only columns with at most 10% gaps. All alignment scoring and filtering procedures were performed using the bppAlnScore and bppSeqMan programs from the Bio++ Program Suite (version 2.4.1) (Guéguen, et al. 2013).

### 4.3 Gene Tree Inference

Gene trees were inferred for each of the 6,012 orthologous groups using IQ-TREE multicore version 2.3.4 (Minh, et al. 2020) under a maximum likelihood framework. Given the large size of the dataset and the need to balance computational efficiency with biological realism, we used the LG4M substitution model for all amino acid alignments.

### 4.4 Detection of Rate Shifts Across Orthologous Groups

To detect evolutionary rate shifts across the 6,012 orthologous groups, we used The RAte Shift EstimatoR (RASER) program (Penn, et al. 2008). For each group, the corresponding protein sequence alignment and maximum likelihood gene tree were provided as input. RASER models sequence evolution under a specified substitution model; in all analyses, we used the default JTT model (Jones, et al. 1992), as the LG model was not supported by the software. For each orthologous group, two independent models were fitted. The null model assumed a uniform evolutionary rate across all branches of the tree, whereas the alternative model allowed for lineage-specific rate shifts along selected branches. Under the alternative model, RASER infers branch-specific rate variation by estimating separate rate parameters while keeping the tree topology fixed. Model fit was assessed using likelihood ratio tests (LRTs), comparing the log-likelihoods of the null and alternative models. Test statistics were evaluated using a chi-squared distribution with three degrees of freedom, following the procedures described in (Penn, et al. 2008), using a significance threshold of α = 0.05. For families showing significant evidence of rate shifts, individual sites were classified as rate-shifting when the posterior probability of the RS model exceeded 0.95. Of the 6,012 orthologous groups analyzed, 4,125 completed successfully under RASER and were retained for all downstream analyses.

### 4.5 Structural mapping and annotation of rate-shifting sites

To investigate structural correlates of evolutionary rate shifts, we integrated site-level rate-shift inference with experimentally resolved protein structures using the SGEDTools (Dutheil, et al. 2024). RASER output files were converted into the Site Group Extended Data (SGED) format using the sged-raser2sged script, providing site-level annotations including amino-acid position and posterior probability of a rate shift. Because RASER analyses were conducted on gap-filtered alignments, site coordinates were translated back to positions in the original unfiltered alignments using the sged-translate-coords script to maintain consistency between evolutionary inference and structural mapping. For each orthologous group, experimentally determined structures were retrieved from the Protein Data Bank (PDB). When multiple candidate structures were available, up to five structures with resolution better than 5 Å were retained, prioritizing those with the highest sequence completeness (i.e., the largest number of modeled residues) using the sged-select-best-structure script. Sequence–structure correspondences were established using sged-create-structure-index applied to the unfiltered MUSCLE5-super5 alignments with a gap penalty of 2 (–gap-index 2). Alignment coordinates were subsequently translated into structural residue coordinates using sged-translate-coords, yielding site-level mappings from alignment positions to PDB residues. Structural descriptors were computed for each mapped site using sged-structure-infos. Extracted features included chain identity, C*α*–C*α* distance distributions among residues (minimum, maximum, mean, and median), relative solvent accessibility as calculated by DSSP (Dictionary of Secondary Structure in Proteins) (Kabsch and Sander 1983), residue depth relative to the molecular surface, and secondary structure classification.

### 4.6 Family-level variation in rate-shifting prevalence

To characterize variation in rate shifts across protein families, we calculated for each of the 4,125 orthologous groups the proportion of alignment sites classified as rate-shifting. We then examined associations between the family-level proportion of rate-shifting sites and two measures of phylogenetic scale: (i) total phylogenetic tree length and (ii) the number of sequences per orthogroup, corresponding to the number of species represented in the family. Associations were assessed using Kendall’s rank correlation coefficient (τ). To evaluate whether the prevalence of rate-shifting sites differs across evolutionary lineages, orthogroups were grouped according to their inferred last common ancestor (LCA), spanning 29 ancestral categories. Differences in rate-shifting proportions among LCA categories were assessed using the Kruskal–Wallis rank-sum test, followed by Dunn’s post hoc tests with correction for multiple comparisons when appropriate.

To disentangle the effects of evolutionary ancestry, taxonomic sampling, and phylogenetic divergence on rate-shift prevalence, we additionally performed residual-based partial correlation analyses. Effects of relevant covariates were first removed using linear models, and Kendall’s rank correlations were subsequently calculated between model residuals. The number of sequences per orthogroup and total phylogenetic tree length were log-transformed prior to residualization. Specifically, we tested whether the association between the proportion of rate-shifting sites and the number of sequences per orthogroup remained significant after correcting for LCA category, and whether the association between rate-shift prevalence and total phylogenetic tree length remained significant after correcting for both LCA category and the number of sequences per orthogroup.

### 4.7 Site-level mixed-effects modeling of rate shifts

To evaluate structural determinants of rate shifts, we modeled the probability that an individual alignment position exhibits a rate shift using generalized linear mixed models with a binomial error distribution and logit link. Analyses were performed on sites that could be mapped to experimentally resolved protein structures. The response variable was binary, indicating whether a site was classified as rate-shifting (1) or non–rate-shifting (0) based on the RASER inference described above. We first evaluated structural effects within a baseline framework that included local structural predictors: relative solvent accessibility (RSA), residue burial (C*α* depth), and secondary structure class (helix, strand, coil). Continuous predictors were standardized to a zero mean and unit variance. Because the ability to detect rate shifts depends on the amount of evolutionary change and taxonomic sampling within a family, we included total phylogenetic tree length and the number of sequences per orthogroup as covariates to control for variation in evolutionary divergence and information content across families. Orthogroup identity was included as a random intercept to account for non-independence among sites within the same protein family. To assess whether baseline rates of rate shifts differ across major evolutionary domains, we extended the baseline model by including evolutionary ancestry as a fixed effect using a four-level classification of last common ancestor (Archaea, Bacteria, Eukaryota, LUCA). Because residue depth was associated with rate shifts in the baseline model, we next introduced an interaction term between C*α* depth and evolutionary ancestry to test whether the effect of burial differs across major evolutionary domains. Finally, to facilitate interpretation of lineage-specific structural effects, we fitted separate mixed models within each ancestry category. Within each lineage-specific subset, continuous predictors were re-standardized to zero mean and unit variance prior to model fitting. These models retained the same structural predictors, phylogenetic covariates, and orthogroup random intercept as the global model, but parameters were estimated independently within each evolutionary domain. Unlike the interaction model, which estimates lineage-specific deviations within a single global framework, the subset approach allows effect sizes to be interpreted relative to the variance structure of each domain separately, albeit with reduced statistical power due to smaller dataset sizes.

All models were fitted using the BinomialBayesMixedGLM class from the statsmodels Python library (Seabold and Perktold 2010), employing variational Bayes estimation for computational scalability. Fixed-effect significance was evaluated using approximate Wald statistics derived from posterior means and standard deviations.

### 4.8 Structural clustering of rate-shifting sites

To assess whether rate-shifting residues are spatially clustered within protein structures, we quantified their three-dimensional proximity using experimentally resolved models. Our approach follows the structural proximity framework of (Chaurasia and Dutheil 2022), adapted here to rate-shifting sites. For each protein family, rate-shifting residues were mapped onto a representative structure using residue-level correspondences established during structural annotation. Only residues with valid C*α* coordinates were retained. Families with fewer than two mapped rate-shifting residues were excluded from structural analyses. Pairwise Euclidean distances between C*α* atoms were computed, and residues separated by less than 8°A were considered in contact. From the resulting contact network, we determined the number of connected components, which reflects the degree of spatial aggregation among rate-shifting sites. This measure was normalized by group size to obtain a dispersion statistic ranging from complete clustering (0) to complete spatial separation (1). One value was computed per protein family. Statistical significance was evaluated using a Monte Carlo permutation test performed independently for each family. For a protein with *m* mapped rate-shifting residues, 1000 random sets of *m* structurally mappable residues were sampled without replacement, preserving the structural geometry of the protein. The same clustering statistic was calculated for each random set, and significance was assessed using a one-sided test to determine whether observed rate-shifting residues were more spatially clustered than expected under random distribution. All structural distances and graph decompositions were computed using the ContactSubgraphs measure implemented in the SgedTools using the sged-structure-infos.py script (Dutheil, et al. 2024).

### 4.9 Effect of rate-shifting sites on phylogenetic reconstruction

To evaluate the impact of rate-shifting (RS) sites on phylogenetic inference, we performed a controlled site-removal analysis on a subset of 50 orthologous protein families. Gene families were randomly sampled from a filtered set of orthogroups that met predefined criteria to ensure sufficient phylogenetic signal (150–500 sequences and at least 300 aligned amino-acid sites) and moderate proportions of rateshifting sites (15–35%). For each gene family, phylogenetic trees were inferred from the full alignment and from resampled alignments. Resampled alignments were generated by removing all sites classified as rate-shifting. To control for the effect of alignment size reduction, 100 replicate alignments per family were generated by randomly removing the same number of sites. In total, this resulted in 5,050 phylogenetic reconstructions (50 full alignments, 50 RS-removed alignments, and 5,000 random replicates). All phylogenetic trees were inferred using IQ-TREE 2 under the LG4M substitution model with ultrafast bootstrap approximation (1,000 replicates) (Minh, et al. 2020). Topological differences between trees inferred from modified alignments and the corresponding full-alignment tree were quantified using normalized Robinson–Foulds (nRF) distances using the ETE v3 framework (Huerta-Cepas, et al. 2016). For each gene family, the effect of RS-site removal was assessed by comparing the nRF distance to the complete-alignment tree obtained after RS-site removal to the distribution of nRF distances obtained from random replicates. Positive differences indicate that removal of RS sites leads to larger topological changes than expected under random site removal. To assess statistical significance at the level of individual gene families, we performed empirical one-sided tests by comparing the observed nRF value after RS-site removal to the distribution of nRF values obtained from random site removal. The p-value was calculated as the proportion of random replicates with equal or greater nRF values, testing whether RS-site removal results in greater topological changes than expected by chance. In addition, to evaluate the effect of RS-site removal on tree support, bootstrap support values were extracted for all internal nodes and averaged per tree. The effect on support was assessed by comparing the mean support after RS-site removal with the distribution of mean support values from random replicates. In addition, to evaluate the effect of RS-site removal on branch length estimation, total phylogenetic tree length was calculated for each inferred tree as the sum of all branch lengths. The effect on branch lengths was assessed by comparing the total tree length obtained after RS-site removal to the distribution of total tree lengths obtained from random replicates. Finally, to test whether variation in the topological impact of RS-site removal across gene families could be explained by alignment characteristics, we fitted linear models with the difference between the nRF distance obtained after RS-site removal and the mean nRF distance across random replicates of the given orthogroup as the response variable, and number of sequences, alignment length, and proportion of rate-shifting sites as predictors.

## Acknowledgements

We acknowledge financial support from the Max Planck Society. M.R.D. is supported by the International Max Planck Research School for Evolutionary Biology (IMPRS EvolBio).

## Author Contributions

M.R.D.: Conceptualization, Data curation, Formal analysis, Investigation, Methodology, Writing – original draft, Writing – review & editing.

E.R.: Formal analysis, Investigation, Writing – review & editing

J.Y.D.: Conceptualization, Methodology, Supervision, Writing – review & editing

## Data Availability Statement

The code used to perform the analyses presented in this study is available at: https://github.com/rdurak/rate-shifts-tree-of-life

## Supplementary Material

Supplementary Table S1. Dunn’s post hoc pairwise comparisons of rate-shifting proportions among orthologous groups classified by inferred last common ancestor (LCA) categories. An overall Kruskal–Wallis rank-sum test was used to test for differences among groups, followed by Dunn’s test for pairwise contrasts. Columns report the two LCA categories compared, the Dunn test statistic (z), the unadjusted p-value, and the Benjamini–Hochberg false discovery rate corrected p-value.

Supplementary Table S2. Baseline mixed-effects logistic regression model predicting the probability that an alignment site is classified as rate-shifting. Continuous predictors were standardized (mean = 0, SD = 1). Reported values include regression coefficients (logit scale), standard errors (SE), Wald z-statistics, p-values, odds ratios (OR), and 95% confidence intervals.

Supplementary Table S3. Mixed-effects logistic regression model including evolutionary ancestry as a fixed effect using four categories (Archaea, Bacteria, Eukaryota, and LUCA), predicting the probability that an alignment site is classified as rate-shifting. Continuous predictors were standardized (mean = 0, SD = 1). Reported values include regression coefficients (logit scale), standard errors (SE), Wald z-statistics, p-values, odds ratios (OR), and 95% confidence intervals.

Supplementary Table S4. Mixed-effects logistic regression model including an interaction between residue burial (Cα depth) and evolutionary ancestry (LCA category), predicting the probability that an alignment site is classified as rate-shifting. Continuous predictors were standardized (mean = 0, SD = 1). Reported values include regression coefficients (logit scale), standard errors (SE), Wald z-statistics, p-values, odds ratios (OR), and 95% confidence intervals.

Supplementary Table S5. Mixed-effects logistic regression model fitted to archaeal orthogroups, predicting the probability that an alignment site is classified as rate-shifting. Continuous predictors were standardized (mean = 0, SD = 1). Reported values include regression coefficients (logit scale), standard errors (SE), Wald z-statistics, p-values, odds ratios (OR), and 95% confidence intervals.

Supplementary Table S6. Mixed-effects logistic regression model fitted to bacterial orthogroups, predicting the probability that an alignment site is classified as rate-shifting. Continuous predictors were standardized (mean = 0, SD = 1). Reported values include regression coefficients (logit scale), standard errors (SE), Wald z-statistics, p-values, odds ratios (OR), and 95% confidence intervals.

Supplementary Table S7. Mixed-effects logistic regression model fitted to eukaryotic orthogroups, predicting the probability that an alignment site is classified as rate-shifting. Continuous predictors were standardized (mean = 0, SD = 1). Reported values include regression coefficients (logit scale), standard errors (SE), Wald z-statistics, p-values, odds ratios (OR), and 95% confidence intervals.

Supplementary Table S8. Mixed-effects logistic regression model fitted to orthogroups inferred to originate from the last universal common ancestor (LUCA), predicting the probability that an alignment site is classified as rate-shifting. Continuous predictors were standardized (mean = 0, SD = 1). Reported values include regression coefficients (logit scale), standard errors (SE), Wald z-statistics, p-values, odds ratios (OR), and 95% confidence intervals.

Supplementary Table S9. Functional annotation of archaeal orthogroups included in the domain-specific structural analysis. The list of 23 orthogroups inferred to originate in Archaea, together with alignment statistics and the proportion of sites classified as rate-shifting.

Supplementary Figure S1. Distribution of the number of protein sequences (species) per orthologous group. Orthogroups contain between 100 and 2,670 sequences, with most families comprising approximately 100–300 sequences.

Supplementary Figure S2. Pairwise differences in rate-shifting proportions across evolutionary categories. Heatmap showing Dunn’s post hoc test z-scores for pairwise comparisons of rate-shifting proportions among orthogroups grouped by their inferred last common ancestor (LCA). Positive z-scores (red) indicate that the row category exhibits a higher median proportion of rate-shifting sites than the column category, whereas negative z-scores (blue) indicate the opposite. For instance, positive values in the LUCA row indicate that LUCA-derived orthogroups exhibit higher proportions of rate-shifting sites than the corresponding column category.

Supplementary Figure S3. Limited spatial clustering of rate-shifting sites. Histogram of p-values from Monte Carlo tests assessing spatial clustering of rate-shifting residues across 1,906 protein families. The red dashed line marks p = 0.05. Only 76 families (4.0%) show significant clustering, indicating that rate-shifting sites are generally dispersed across protein structures.

