## Supplementary Table 1 for "The Shifting Tempo of Evolution: Mapping Site-Specific Rate Shifts Across the Tree of Life"

Supplementary Table S1. Dunn’s post hoc pairwise comparisons of rate-shifting proportions among orthologous groups classified by inferred last common ancestor (LCA) categories. An overall Kruskal–Wallis rank-sum test was used to test for differences among groups, followed by Dunn’s test for pairwise contrasts. Columns report the two LCA categories compared, the Dunn test statistic (z), the unadjusted p-value, and the Benjamini–Hochberg false discovery rate corrected p-value.

| LCA Group 1 | LCA Group 2 | Z statistic | Raw p value | FDR-adjusted p-value |
| --- | --- | --- | --- | --- |
| Bilateria | LUCA | -12.698660803908837 | 0.0 | 0.0 |
| Gnathostomata | LUCA | -16.608568072513307 | 0.0 | 0.0 |
| Eukaryota | Gnathostomata | 12.873175384213472 | 0.0 | 0.0 |
| LUCA | Vertebrata | 10.308949179083173 | 0.0 | 0.0 |
| Euteleostomi | LUCA | -8.950333805073424 | 0.0 | 0.0 |
| Eumetazoa | LUCA | -10.991958299607253 | 0.0 | 0.0 |
| Eukaryota | Vertebrata | 8.368780666271954 | 1.1102230246251565e-16 | 1.6653345369377348e-15 |
| Bilateria | Eukaryota | -8.35366707200308 | 1.1102230246251565e-16 | 1.6653345369377348e-15 |
| Gnathostomata | Opisthokonta | -8.104724675035646 | 5.551115123125783e-16 | 7.401486830834377e-15 |
| Bacteria | LUCA | -7.714217586069573 | 1.2212453270876722e-14 | 1.4654943925052066e-13 |
| Opisthokonta | Vertebrata | 7.165890704187971 | 7.728262474415715e-13 | 8.43083179027169e-12 |
| Eukaryota | Eumetazoa | 7.137161403596667 | 9.527933997333093e-13 | 9.527933997333093e-12 |
| LUCA | Metazoa | 6.947513517444499 | 3.717803842562262e-12 | 3.431818931595934e-11 |
| Chordata | LUCA | -6.868018203583579 | 6.5100147494945304e-12 | 5.5800126424238834e-11 |
| Fungi | Vertebrata | 6.778428449043024 | 1.2149059536170626e-11 | 9.7192476289365e-11 |
| Fungi | Gnathostomata | 6.5658197031488506 | 5.1747384155476084e-11 | 3.8810538116607063e-10 |
| Bacteria | Gnathostomata | 6.275359439086564 | 3.488275224228232e-10 | 2.4623119229846346e-09 |
| Eukaryota | LUCA | -6.160856471858549 | 7.235254617654618e-10 | 4.8235030784364126e-09 |
| Eukaryota | Euteleostomi | 6.076251312872307 | 1.230245572081401e-09 | 7.769972034198322e-09 |
| Bacteria | Vertebrata | 5.820695828462546 | 5.860313567751518e-09 | 3.516188140650911e-08 |
| Chordata | Eukaryota | -5.571550903600348 | 2.5248157009549743e-08 | 1.4427518291171282e-07 |
| Chordata | Fungi | -5.292517861674024 | 1.206436821554746e-07 | 6.580564481207706e-07 |
| Deuterostomia | LUCA | -5.22009693367471 | 1.7882950609227066e-07 | 9.330235100466295e-07 |
| Chordata | Opisthokonta | -5.109285932897914 | 3.233786474154954e-07 | 1.616893237077477e-06 |
| Dikarya | Vertebrata | 4.95295788354658 | 7.309379946862649e-07 | 3.5085023744940712e-06 |
| Bilateria | Vertebrata | 4.782878055736037 | 1.7280296670252326e-06 | 7.975521540116458e-06 |
| Bilateria | Gnathostomata | 4.761101977272589 | 1.9253872098889957e-06 | 8.557276488395536e-06 |
| Bilateria | Opisthokonta | -4.708786287231143 | 2.4919617813701223e-06 | 1.0679836205871953e-05 |
| Eumetazoa | Vertebrata | 4.660550062389981 | 3.1536544126131716e-06 | 1.3049604465985537e-05 |
| Eukaryota | Metazoa | 4.591197155220894 | 4.407107760950879e-06 | 1.7628431043803516e-05 |
| Euteleostomi | Opisthokonta | -4.45325704448141 | 8.457740037215444e-06 | 3.2739638853737204e-05 |
| Archaea | LUCA | -4.444592230598725 | 8.805872561024053e-06 | 3.30220221038402e-05 |
| Eumetazoa | Opisthokonta | -4.4100773373352995 | 1.0333370830228894e-05 | 3.757589392810507e-05 |
| Euteleostomi | Fungi | -4.345699385000053 | 1.3883259713520779e-05 | 4.8077104192602504e-05 |
| Eumetazoa | Gnathostomata | 4.343508754050374 | 1.4022488722842397e-05 | 4.8077104192602504e-05 |
| Chordata | Dikarya | -4.281658218866741 | 1.855057543509453e-05 | 6.18352514503151e-05 |
| Bilateria | Fungi | -4.254979506532006 | 2.090681630029234e-05 | 6.780589070365084e-05 |
| Dikarya | Gnathostomata | 4.170107203465549 | 3.044563871368311e-05 | 9.614412225373614e-05 |
| Eumetazoa | Fungi | -4.134543583840181 | 3.556607554999758e-05 | 0.00010943407861537716 |
| Bacteria | Chordata | 4.077941062818213 | 4.5436280012411956e-05 | 0.00013630884003723587 |
| Bacteria | Eukaryota | -4.014412082607244 | 5.9594151316066934e-05 | 0.0001744219062909276 |
| Deuterostomia | Eukaryota | -3.8665118509420693 | 0.00011040310142063081 | 0.00031543743263037375 |
| Deuterostomia | Fungi | -3.827168451200456 | 0.00012962580922326605 | 0.0003617464443439983 |
| Fungi | Metazoa | 3.815639247765708 | 0.00013583083032819854 | 0.0003704477190769051 |
| Euteleostomi | Vertebrata | 3.714995713580724 | 0.00020320713905730425 | 0.0005418857041528113 |
| LUCA | Opisthokonta | 3.634795857923148 | 0.0002782008362435251 | 0.0007257413119396307 |
| Metazoa | Opisthokonta | -3.6111088519334036 | 0.000304890653699319 | 0.0007784442222110272 |
| Metazoa | Vertebrata | 3.6050325847012092 | 0.0003121137652725148 | 0.0007802844131812869 |
| Deuterostomia | Opisthokonta | -3.4772408290466807 | 0.0005066025579260058 | 0.0012406593255330756 |
| Bilateria | Chordata | 3.2960754343864154 | 0.000980456852114786 | 0.0023530964450754867 |
| Archaea | Fungi | -3.2619141862310324 | 0.0011066263789792918 | 0.002567044896376175 |
| Chordata | Eumetazoa | -3.260442263202353 | 0.0011123861217630093 | 0.002567044896376175 |
| Archaea | Eukaryota | -3.1662377415570027 | 0.0015442452543137364 | 0.0034964043493895917 |
| Deuterostomia | Dikarya | -3.04355234149858 | 0.002338027649453789 | 0.005195616998786197 |
| Dikarya | Euteleostomi | 2.8740252982583874 | 0.004052764987934054 | 0.008842396337310664 |
| Archaea | Opisthokonta | -2.8465672461102605 | 0.004419339418024193 | 0.009405611025546993 |
| Bacteria | Fungi | -2.8431028383060055 | 0.004467665237134821 | 0.009405611025546993 |
| Chordata | Metazoa | -2.755560793371767 | 0.005859161500707755 | 0.012122403104912596 |
| Chordata | Euteleostomi | -2.7339802196210683 | 0.006257380334541263 | 0.012726875256694093 |
| Bilateria | Dikarya | -2.627931674983506 | 0.008590576839529684 | 0.016944198284136843 |
| Dikarya | Metazoa | 2.627032891307331 | 0.00861330079443623 | 0.016944198284136843 |
| Archaea | Dikarya | -2.621344513017876 | 0.008758370432133344 | 0.01695168470735486 |
| Dikarya | Eumetazoa | 2.5936458580968536 | 0.009496424860483077 | 0.01780579661340577 |
| Ascomycota | Vertebrata | 2.597267513969783 | 0.009396870811599567 | 0.01780579661340577 |
| Euteleostomi | Gnathostomata | 2.5579909413135486 | 0.010527883469942156 | 0.01943609255989321 |
| Bacteria | Euteleostomi | 2.525711784847997 | 0.011546416355024514 | 0.02099348428186275 |
| Bacteria | Opisthokonta | -2.4511170214291207 | 0.014241364566459302 | 0.0255069216115689 |
| Gnathostomata | Metazoa | -2.399929929131297 | 0.016398210521481604 | 0.028938018567320478 |
| Ascomycota | Chordata | 2.3699625488995992 | 0.017789887156089534 | 0.030496949410439202 |
| Bacteria | Deuterostomia | 2.3737721925050193 | 0.017607410007054125 | 0.030496949410439202 |
| Ascomycota | LUCA | -2.315628650787851 | 0.02057855312067891 | 0.034780653161710834 |
| Bacteria | Bilateria | 2.307879484163099 | 0.021005840440384005 | 0.035009734067306675 |
| Gnathostomata | Vertebrata | 2.256692421090373 | 0.024027300756467862 | 0.03949693275035813 |
| Bacteria | Eumetazoa | 2.114018670991281 | 0.03451367265709693 | 0.05596811782231934 |
| Bacteria | Metazoa | 1.8815253810301662 | 0.059900485996489006 | 0.0958407775943824 |
| Archaea | Vertebrata | 1.8604974341000202 | 0.06281518114399387 | 0.09918186496420085 |
| Archaea | Bacteria | -1.785849355685927 | 0.07412365285586231 | 0.11551738107407113 |
| Bacteria | Dikarya | -1.7671348425689817 | 0.07720564856356449 | 0.1187779208670223 |
| Ascomycota | Gnathostomata | 1.6553181494799378 | 0.0978599792655499 | 0.14778803146064412 |
| Archaea | Chordata | 1.652045114694518 | 0.09852535430709608 | 0.14778803146064412 |
| Ascomycota | Fungi | -1.645520402050438 | 0.09986253885041296 | 0.1479445020006118 |
| Chordata | Gnathostomata | -1.6321534860489288 | 0.10264715665891866 | 0.15021535120817364 |
| Bilateria | Deuterostomia | 1.5281745584567388 | 0.12646920372192916 | 0.1828470415256807 |
| Deuterostomia | Vertebrata | 1.5190337005371015 | 0.1287540133823445 | 0.18393430483192072 |
| Deuterostomia | Eumetazoa | -1.5129471461631288 | 0.1302930848735434 | 0.18394317864500248 |
| Ascomycota | Dikarya | -1.3653184475469278 | 0.17215298335856677 | 0.24021346515148853 |
| Chordata | Deuterostomia | -1.3499263367785852 | 0.1770396127548105 | 0.24419256931698 |
| Ascomycota | Eukaryota | -1.3076561457327647 | 0.19098996412029545 | 0.2604408601640392 |
| Ascomycota | Deuterostomia | 1.251557038821158 | 0.21073131728225358 | 0.2841321131895554 |
| Ascomycota | Opisthokonta | -1.1429086303960228 | 0.25307652890627086 | 0.3374353718750278 |
| Deuterostomia | Metazoa | -1.1349774682197684 | 0.2563847591299939 | 0.3380897922593326 |
| Fungi | Opisthokonta | 1.1279788804735766 | 0.2593288385220943 | 0.3382550067679491 |
| Deuterostomia | Euteleostomi | -1.03995836930314 | 0.2983592424530366 | 0.3849796676813375 |
| Eukaryota | Fungi | -1.0279700272323948 | 0.30396392693584584 | 0.3880390556627819 |
| Archaea | Eumetazoa | -0.9673057656060623 | 0.33339120230013064 | 0.41244272449500696 |
| Archaea | Bilateria | -0.9741000713511163 | 0.3300068541108241 | 0.41244272449500696 |
| Fungi | LUCA | -0.972695823997579 | 0.3307045013643466 | 0.41244272449500696 |
| Archaea | Ascomycota | -0.9291119850494317 | 0.35283105392952796 | 0.4320380252198302 |
| Bilateria | Euteleostomi | 0.8868280938941743 | 0.3751714597179472 | 0.45475328450660263 |
| Eumetazoa | Euteleostomi | 0.8490683451653086 | 0.3958432632435237 | 0.4750119158922284 |
| Dikarya | LUCA | -0.7272531624908917 | 0.4670708846561934 | 0.554935704542012 |
| Ascomycota | Euteleostomi | 0.6871563330163487 | 0.49198421834337247 | 0.5788049627569088 |
| Dikarya | Opisthokonta | 0.6565032115389303 | 0.5115004018733547 | 0.5903008116449397 |
| Archaea | Metazoa | -0.656357643320846 | 0.5115940367589478 | 0.5903008116449397 |
| Archaea | Gnathostomata | 0.6343259601445334 | 0.5258681179233728 | 0.6009921347695689 |
| Ascomycota | Metazoa | 0.5582033820604972 | 0.5767055116795146 | 0.6528741641654882 |
| Archaea | Euteleostomi | -0.5431787983399871 | 0.5870067027848374 | 0.6578536300904534 |
| Dikarya | Eukaryota | 0.5358412552145912 | 0.5920682670814081 | 0.6578536300904534 |
| Bilateria | Metazoa | 0.46011521883318257 | 0.6454335208513191 | 0.7105690137812687 |
| Eumetazoa | Metazoa | 0.45118188319732133 | 0.6518584658363576 | 0.7111183263669357 |
| Eukaryota | Opisthokonta | 0.4077796258513168 | 0.683435472858398 | 0.7388491598469167 |
| Ascomycota | Eumetazoa | 0.38703040784815074 | 0.6987336966167522 | 0.7420180849027457 |
| Ascomycota | Bilateria | 0.3936309535677406 | 0.6938535248425306 | 0.7420180849027457 |
| Archaea | Deuterostomia | 0.3491253162329226 | 0.726995231335714 | 0.76525813824812 |
| Ascomycota | Bacteria | -0.26946681864063404 | 0.7875704730324566 | 0.8218126675121287 |
| Euteleostomi | Metazoa | -0.23182853656512117 | 0.8166711918004945 | 0.8448322673798219 |
| Deuterostomia | Gnathostomata | 0.18114453447782977 | 0.8562541293118948 | 0.8782093633968152 |
| Chordata | Vertebrata | -0.0972702025008908 | 0.9225118187243354 | 0.9314186114047546 |
| Dikarya | Fungi | -0.09582849011386198 | 0.9236567896430483 | 0.9314186114047546 |
| Bilateria | Eumetazoa | -0.013699903539200739 | 0.9890694004052107 | 0.9890694004052107 |
