## Supplementary Table 2 for "The Shifting Tempo of Evolution: Mapping Site-Specific Rate Shifts Across the Tree of Life"

Supplementary Table S2. Baseline mixed-effects logistic regression model predicting the probability that an alignment site is classified as rate-shifting. Continuous predictors were standardized (mean = 0, SD = 1). Reported values include regression coefficients (logit scale), standard errors (SE), Wald z-statistics, p-values, odds ratios (OR), and 95% confidence intervals.

| Predictor | Logit coefficient | SE | z-statistic | p_value | Odds ratio | 95% CI (lower) | 95% CI (upper) |
| --- | --- | --- | --- | --- | --- | --- | --- |
| Intercept | -1.9220 | 0.0060 | -318.2836 | 0.0000 | 0.1463 | 0.1445 | 0.1480 |
| C(SS3)[T.Helix] | -0.0018 | 0.0099 | -0.1823 | 0.8553 | 0.9981 | 0.9788 | 1.0178 |
| C(SS3)[T.Strand] | -0.0144 | 0.0128 | -1.1252 | 0.2604 | 0.9856 | 0.9611 | 1.0107 |
| Rsa_z | -0.0001 | 0.0060 | -0.0138 | 0.9890 | 0.9999 | 0.9882 | 1.0118 |
| Depth_z | -0.0213 | 0.0062 | -3.4704 | 0.0005 | 0.9789 | 0.9671 | 0.9908 |
| log_tree_z | 0.6348 | 0.0061 | 103.4109 | 0.0000 | 1.8867 | 1.8642 | 1.9096 |
| log_numseq_z | 0.2237 | 0.0051 | 43.7819 | 0.0000 | 1.2506 | 1.2382 | 1.2632 |
