## Supplementary Table 3 for "The Shifting Tempo of Evolution: Mapping Site-Specific Rate Shifts Across the Tree of Life"

Supplementary Table S3. Mixed-effects logistic regression model including evolutionary ancestry as a fixed effect using four categories (Archaea, Bacteria, Eukaryota, and LUCA), predicting the probability that an alignment site is classified as rate-shifting. Continuous predictors were standardized (mean = 0, SD = 1). Reported values include regression coefficients (logit scale), standard errors (SE), Wald z-statistics, p-values, odds ratios (OR), and 95% confidence intervals.

| Predictor | Logit coefficient | SE | z-statistic | p_value | Odds ratio | 95% CI (lower) | 95% CI (upper) |
| --- | --- | --- | --- | --- | --- | --- | --- |
| Intercept | -2.8925 | 0.0060 | -478.9748 | 0.0000 | 0.0554 | 0.0548 | 0.0561 |
| C(LCA_2)[T.Bacteria] | 0.2815 | 0.0186 | 15.1280 | 0.0000 | 1.3251 | 1.2776 | 1.3743 |
| C(LCA_2)[T.Eukaryota] | 1.1120 | 0.0081 | 136.9534 | 0.0000 | 3.0406 | 2.9926 | 3.0893 |
| C(LCA_2)[T.LUCA] | 0.8264 | 0.0104 | 79.2273 | 0.0000 | 2.2852 | 2.2389 | 2.3324 |
| C(SS3)[T.Helix] | -0.0016 | 0.0100 | -0.1592 | 0.8735 | 0.9984 | 0.9791 | 1.0181 |
| C(SS3)[T.Strand] | -0.0144 | 0.0128 | -1.1193 | 0.2630 | 0.9857 | 0.9613 | 1.0108 |
| Rsa_z | -0.0001 | 0.0060 | -0.0205 | 0.9836 | 0.9999 | 0.9881 | 1.0118 |
| Depth_z | -0.0205 | 0.0062 | -3.3274 | 0.0009 | 0.9797 | 0.9680 | 0.9916 |
| log_tree_z | 0.7745 | 0.0061 | 126.1463 | 0.0000 | 2.1696 | 2.1436 | 2.1958 |
| log_numseq_z | 0.1784 | 0.0051 | 34.9151 | 0.0000 | 1.1953 | 1.1834 | 1.2073 |
