## Supplementary Table 4 for "The Shifting Tempo of Evolution: Mapping Site-Specific Rate Shifts Across the Tree of Life"

Supplementary Table S4. Mixed-effects logistic regression model including an interaction between residue burial (Cα depth) and evolutionary ancestry (LCA category), predicting the probability that an alignment site is classified as rate-shifting. Continuous predictors were standardized (mean = 0, SD = 1). Reported values include regression coefficients (logit scale), standard errors (SE), Wald z-statistics, p-values, odds ratios (OR), and 95% confidence intervals.

| Predictor | Logit coefficient | SE | z-statistic | p_value | Odds ratio | 95% CI (lower) | 95% CI (upper) |
| --- | --- | --- | --- | --- | --- | --- | --- |
| Intercept | -2.8673 | 0.0060 | -474.7992 | 0.0000 | 0.0569 | 0.0562 | 0.0575 |
| C(LCA_2)[T.Bacteria] | 0.2568 | 0.0186 | 13.8007 | 0.0000 | 1.2928 | 1.2465 | 1.3408 |
| C(LCA_2)[T.Eukaryota] | 1.0864 | 0.0081 | 133.7859 | 0.0000 | 2.9635 | 2.9167 | 3.0111 |
| C(LCA_2)[T.LUCA] | 0.8009 | 0.0104 | 76.7732 | 0.0000 | 2.2274 | 2.1824 | 2.2734 |
| C(SS3)[T.Helix] | -0.0015 | 0.0100 | -0.1492 | 0.8814 | 0.9985 | 0.9792 | 1.0182 |
| C(SS3)[T.Strand] | -0.0146 | 0.0128 | -1.1366 | 0.2557 | 0.9855 | 0.9610 | 1.0106 |
| Rsa_z | 0.0002 | 0.0060 | 0.0367 | 0.9707 | 1.0002 | 0.9885 | 1.0121 |
| Depth_z | 0.3788 | 0.0062 | 61.3798 | 0.0000 | 1.4605 | 1.4429 | 1.4782 |
| C(LCA_2)[T.Bacteria]:Depth_z | -0.4271 | 0.0220 | -19.3927 | 0.0000 | 0.6524 | 0.6248 | 0.6812 |
| C(LCA_2)[T.Eukaryota]:Depth_z | -0.4049 | 0.0104 | -38.8719 | 0.0000 | 0.6671 | 0.6536 | 0.6808 |
| C(LCA_2)[T.LUCA]:Depth_z | -0.3849 | 0.0082 | -47.0909 | 0.0000 | 0.6805 | 0.6697 | 0.6915 |
| log_tree_z | 0.7747 | 0.0061 | 126.1837 | 0.0000 | 2.1700 | 2.1440 | 2.1963 |
| log_numseq_z | 0.1782 | 0.0051 | 34.8806 | 0.0000 | 1.1951 | 1.1832 | 1.2071 |
