## Supplementary Table 5 for "The Shifting Tempo of Evolution: Mapping Site-Specific Rate Shifts Across the Tree of Life"

Supplementary Table S5. Mixed-effects logistic regression model fitted to archaeal orthogroups, predicting the probability that an alignment site is classified as rate-shifting. Continuous predictors were standardized (mean = 0, SD = 1). Reported values include regression coefficients (logit scale), standard errors (SE), Wald z-statistics, p-values, odds ratios (OR), and 95% confidence intervals.

| Predictor | Logit coefficient | SE | z-statistic | p_value | Odds ratio | 95% CI (lower) | 95% CI (upper) |
| --- | --- | --- | --- | --- | --- | --- | --- |
| Intercept | -2.2793 | 0.0751 | -30.3446 | 0.0000 | 0.1024 | 0.0883 | 0.1186 |
| C(SS3)[T.Helix] | -0.1156 | 0.1186 | -0.9748 | 0.3297 | 0.8908 | 0.7061 | 1.1239 |
| C(SS3)[T.Strand] | -0.3320 | 0.1593 | -2.0841 | 0.0372 | 0.7175 | 0.5251 | 0.9804 |
| Rsa_z | 0.0753 | 0.0724 | 1.0403 | 0.2982 | 1.0782 | 0.9356 | 1.2426 |
| Depth_z | 0.1998 | 0.0711 | 2.8117 | 0.0049 | 1.2211 | 1.0624 | 1.4036 |
| log_tree_z | 0.2441 | 0.0730 | 3.3436 | 0.0008 | 1.2765 | 1.1063 | 1.4729 |
| log_numseq_z | 0.5996 | 0.0732 | 8.1889 | 0.0000 | 1.8214 | 1.5779 | 2.1025 |
