## Supplementary Table 6 for "The Shifting Tempo of Evolution: Mapping Site-Specific Rate Shifts Across the Tree of Life"

Supplementary Table S6. Mixed-effects logistic regression model fitted to bacterial orthogroups, predicting the probability that an alignment site is classified as rate-shifting. Continuous predictors were standardized (mean = 0, SD = 1). Reported values include regression coefficients (logit scale), standard errors (SE), Wald z-statistics, p-values, odds ratios (OR), and 95% confidence intervals.

| Predictor | Logit coefficient | SE | z-statistic | p_value | Odds ratio | 95% CI (lower) | 95% CI (upper) |
| --- | --- | --- | --- | --- | --- | --- | --- |
| Intercept | -1.9364 | 0.0185 | -104.5992 | 0.0000 | 0.1442 | 0.1391 | 0.1495 |
| C(SS3)[T.Helix] | -0.0027 | 0.0280 | -0.0948 | 0.9244 | 0.9973 | 0.9441 | 1.0536 |
| C(SS3)[T.Strand] | -0.0642 | 0.0416 | -1.5455 | 0.1222 | 0.9378 | 0.8644 | 1.0174 |
| Rsa_z | 0.0113 | 0.0184 | 0.6113 | 0.5410 | 1.0113 | 0.9754 | 1.0486 |
| Depth_z | -0.0177 | 0.0177 | -0.9995 | 0.3175 | 0.9825 | 0.9491 | 1.0171 |
| log_tree_z | 0.3559 | 0.0193 | 18.4426 | 0.0000 | 1.4274 | 1.3745 | 1.4825 |
| log_numseq_z | 0.4034 | 0.0168 | 23.9701 | 0.0000 | 1.4969 | 1.4484 | 1.5471 |
