## Supplementary Table 7 for "The Shifting Tempo of Evolution: Mapping Site-Specific Rate Shifts Across the Tree of Life"

Supplementary Table S7. Mixed-effects logistic regression model fitted to eukaryotic orthogroups, predicting the probability that an alignment site is classified as rate-shifting. Continuous predictors were standardized (mean = 0, SD = 1). Reported values include regression coefficients (logit scale), standard errors (SE), Wald z-statistics, p-values, odds ratios (OR), and 95% confidence intervals.

| Predictor | Logit coefficient | SE | z-statistic | p_value | Odds ratio | 95% CI (lower) | 95% CI (upper) |
| --- | --- | --- | --- | --- | --- | --- | --- |
| Intercept | -2.2450 | 0.0082 | -275.2327 | 0.0000 | 0.1059 | 0.1042 | 0.1076 |
| C(SS3)[T.Helix] | 0.0010 | 0.0139 | 0.0722 | 0.9424 | 1.0010 | 0.9742 | 1.0285 |
| C(SS3)[T.Strand] | -0.0283 | 0.0170 | -1.6601 | 0.0969 | 0.9721 | 0.9402 | 1.0051 |
| Rsa_z | -0.0083 | 0.0081 | -1.0213 | 0.3071 | 0.9917 | 0.9761 | 1.0077 |
| Depth_z | -0.0284 | 0.0090 | -3.1450 | 0.0017 | 0.9720 | 0.9549 | 0.9893 |
| log_tree_z | 0.8772 | 0.0080 | 109.6010 | 0.0000 | 2.4042 | 2.3668 | 2.4422 |
| log_numseq_z | 0.0683 | 0.0067 | 10.2540 | 0.0000 | 1.0707 | 1.0568 | 1.0848 |
