## Supplementary Table 8 for "The Shifting Tempo of Evolution: Mapping Site-Specific Rate Shifts Across the Tree of Life"

Supplementary Table S8. Mixed-effects logistic regression model fitted to orthogroups inferred to originate from the last universal common ancestor (LUCA), predicting the probability that an alignment site is classified as rate-shifting. Continuous predictors were standardized (mean = 0, SD = 1). Reported values include regression coefficients (logit scale), standard errors (SE), Wald z-statistics, p-values, odds ratios (OR), and 95% confidence intervals.

| Predictor | Logit coefficient | SE | z-statistic | p_value | Odds ratio | 95% CI (lower) | 95% CI (upper) |
| --- | --- | --- | --- | --- | --- | --- | --- |
| Intercept | -1.0974 | 0.0104 | -105.9624 | 0.0000 | 0.3337 | 0.3270 | 0.3406 |
| C(SS3)[T.Helix] | -0.0030 | 0.0168 | -0.1780 | 0.8587 | 0.9970 | 0.9647 | 1.0304 |
| C(SS3)[T.Strand] | 0.0182 | 0.0224 | 0.8133 | 0.4161 | 1.0183 | 0.9747 | 1.0640 |
| Rsa_z | 0.0108 | 0.0103 | 1.0404 | 0.2982 | 1.0108 | 0.9905 | 1.0315 |
| Depth_z | 0.0139 | 0.0109 | 1.2745 | 0.2025 | 1.0140 | 0.9926 | 1.0359 |
| log_tree_z | 0.0801 | 0.0111 | 7.2307 | 0.0000 | 1.0834 | 1.0601 | 1.1072 |
| log_numseq_z | 0.3869 | 0.0104 | 37.2560 | 0.0000 | 1.4725 | 1.4428 | 1.5027 |
