## Supplementary Table 9 for "The Shifting Tempo of Evolution: Mapping Site-Specific Rate Shifts Across the Tree of Life"

Supplementary Table S9. Functional annotation of archaeal orthogroups included in the domain-specific structural analysis. The list of 23 orthogroups inferred to originate in Archaea, together with alignment statistics and the proportion of sites classified as rate-shifting.

| Orthogroup ID | Alignment sites | Number of rate-shifting sites | Rate-shift proportion | Inferred last common ancestor | Functional annotation |
| --- | --- | --- | --- | --- | --- |
| AIYDHKT | 152 | 15 | 0.098 | Archaea | protein of unknown function DUF359 |
| CRECHRI | 58 | 3 | 0.051 | Archaea | ÊDNA-directed RNA polymerase subunit |
| DIVPCYK | 385 | 68 | 0.176 | Archaea | CCA-adding enzyme |
| DSWIEIQ | 89 | 14 | 0.157 | Archaea | translation initiation factor |
| DTGYAYH | 203 | 32 | 0.157 | Archaea | phosphoglycolate phosphatase |
| DWTEKYR | 380 | 82 | 0.215 | Archaea | AAA ATPase central domain protein |
| EAYVYPT | 122 | 8 | 0.065 | Archaea | hypothetical protein |
| EMSHPGA | 190 | 24 | 0.126 | Archaea | orotidine 5'-phosphate decarboxylase |
| FADMRMA | 495 | 117 | 0.236 | Archaea | PilT protein domain protein |
| FSEIEHF | 142 | 53 | 0.373 | Archaea | transcription elongation factor Spt5 |
| GNTLEHW | 197 | 12 | 0.060 | Archaea | DNA repair and recombination protein radB |
| GYDVHMN | 69 | 6 | 0.086 | Archaea | putative snRNP Sm-like protein |
| GYHVHVR | 274 | 41 | 0.149 | Archaea | DNA primase small subunit |
| HAEYEDY | 180 | 12 | 0.066 | Archaea | riboflavin kinase |
| KGMPHRR | 93 | 14 | 0.150 | Archaea | ribosomal protein L21e |
| KRRNCDG | 1050 | 71 | 0.067 | Archaea | DNA polymerase II large subunit |
| MLGETVF | 227 | 26 | 0.114 | Archaea | GHMP kinase |
| PRKMKQM | 105 | 7 | 0.066 | Archaea | nascent polypeptide-associated complex protein |
| QYIRSYF | 97 | 0 | 0 | Archaea | Êribosomal protein S24e |
| RTIIWEV | 81 | 2 | 0.024 | Archaea | hypothetical protein |
| SGVIIME | 110 | 23 | 0.209 | Archaea | 50S ribosomal protein L18e |
| WIARHAK | 79 | 16 | 0.202 | Archaea | ribosomal protein L31e |
| YAMHSNA | 179 | 0 | 0 | Archaea | adenylate kinase |
