## Supplementary figures and images for "The Shifting Tempo of Evolution: Mapping Site-Specific Rate Shifts Across the Tree of Life"

### Supplementary Figure 1

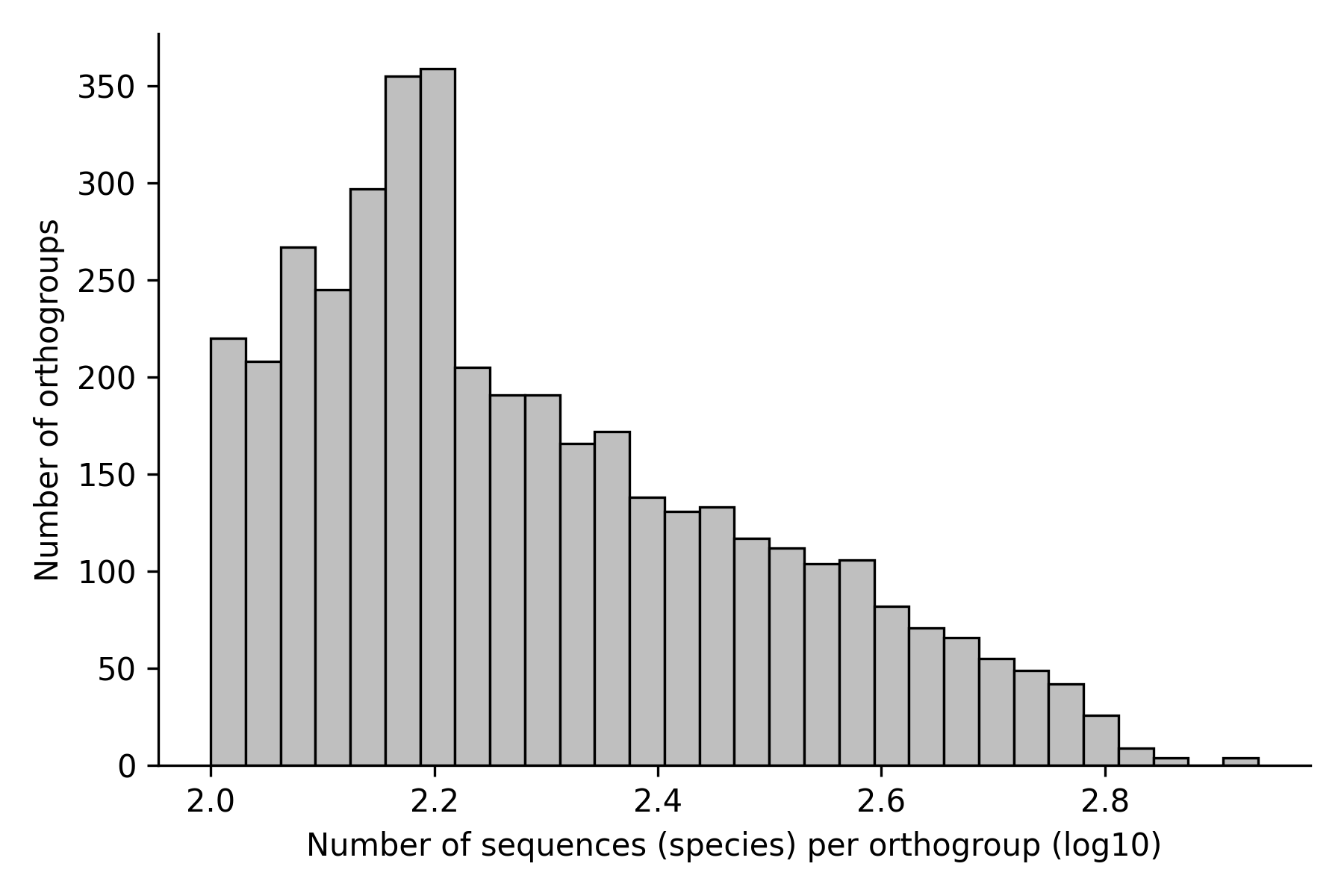

### Supplementary Figure 2

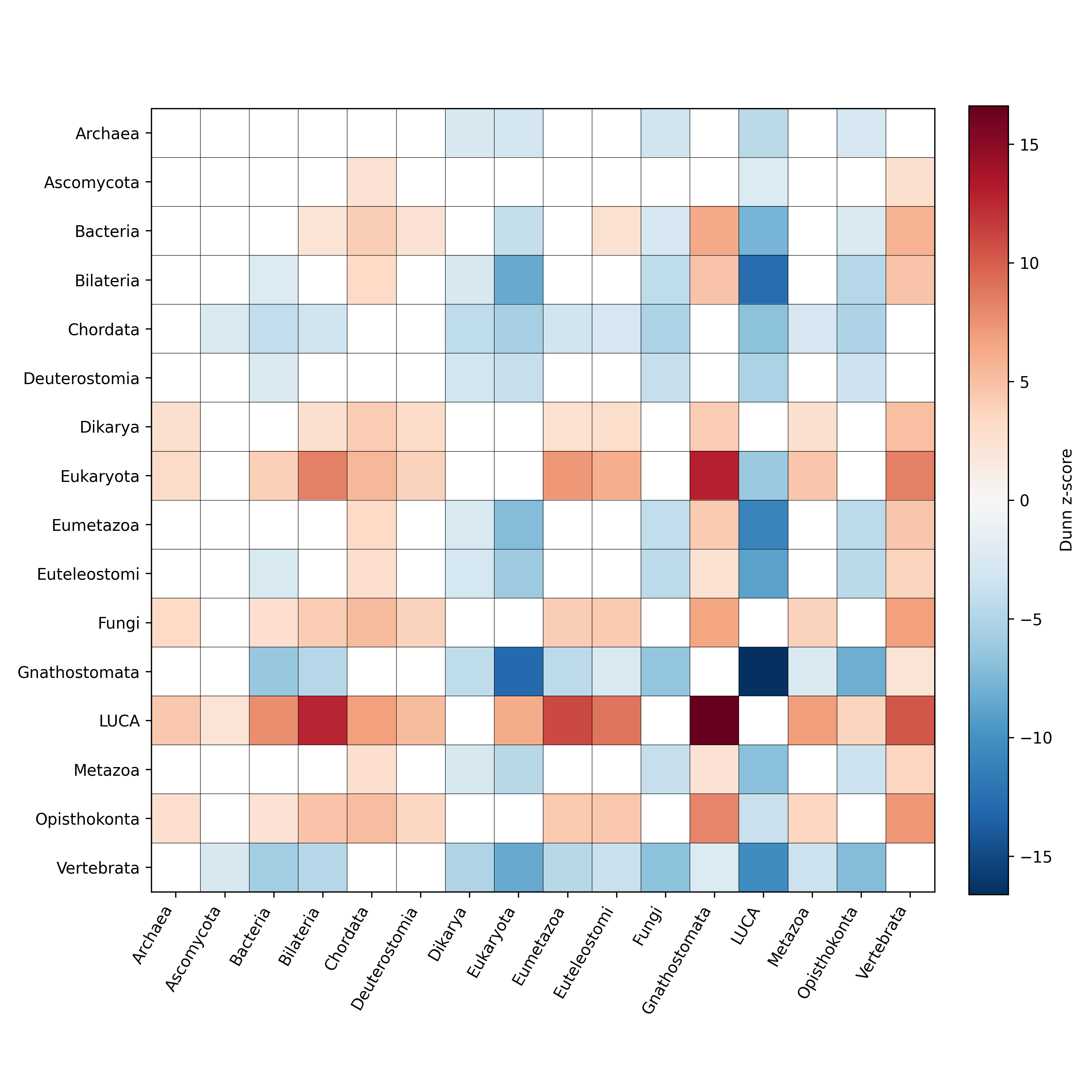

### Supplementary Figure 3

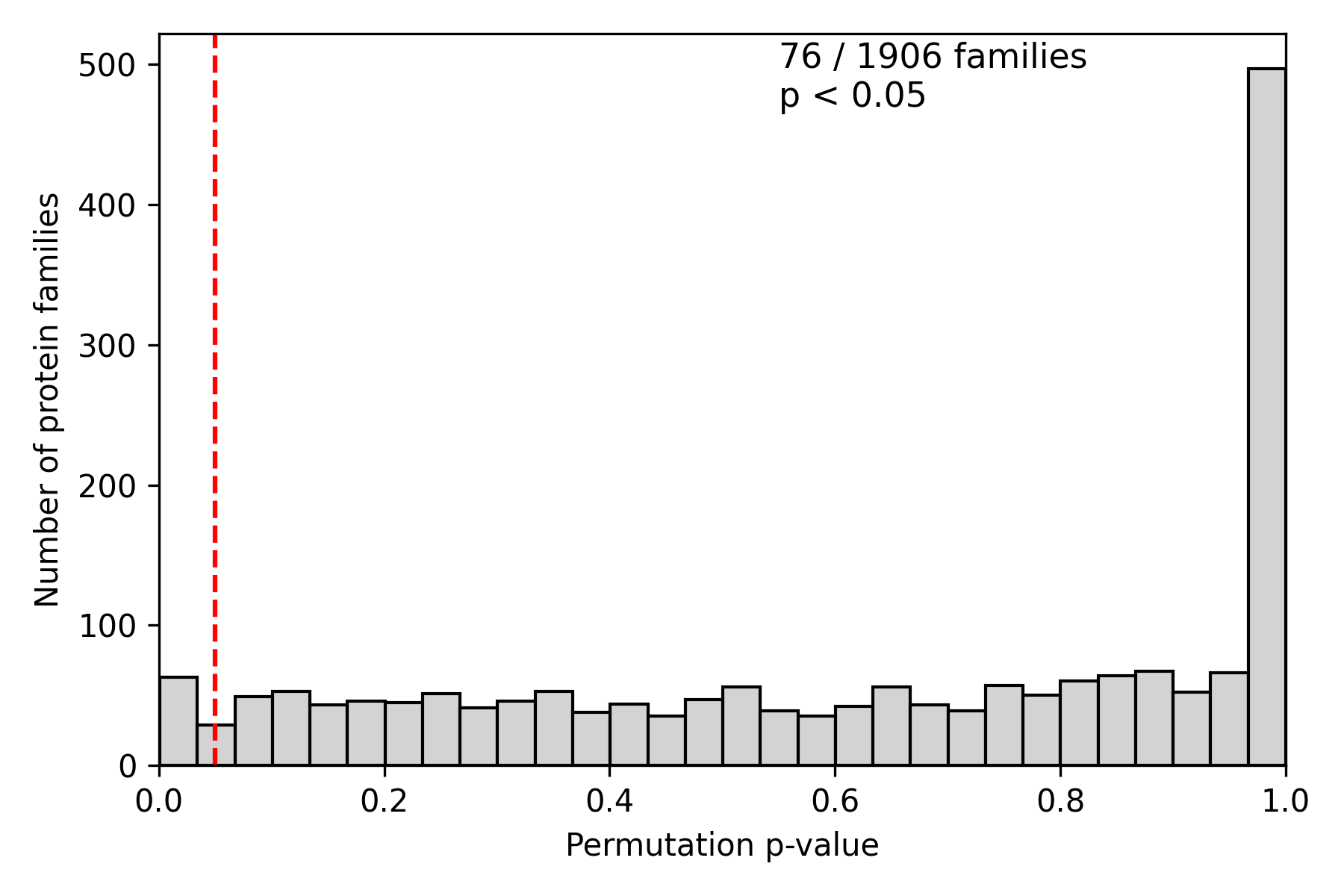
